# Anoxygenic phototrophic Chlorobi use broad metabolic and resource acquisition strategies to support stable near-clonal blooms

**DOI:** 10.64898/2026.09.24.754110

**Authors:** Molly A. Moynihan, Olivia L. Mathieson, Catherine A. Crowley, Eleanor Greene, Hannah Vanderscheuren, Diana Dumit, Roo Weed, Ketil Koop-Jakobsen, Manuel Kleiner, S. Emil Ruff

## Abstract

Anoxygenic phototrophic green sulfur bacteria (GSB; Chlorobiia) are important primary producers in anoxic and sulfidic environments, whose significance in aquatic ecosystems will expand with coastal deoxygenation. Here, we employed a uniquely comprehensive analytical approach to investigate the formation, maintenance, and collapse of a GSB bloom. We combined multi-omics—V4V5 and synthetic long read 16S rRNA amplicon sequencing, metagenomics, and metaproteomics—with total and GSB-specific cell counts, biogeochemical measurements, and isotopic analysis. The GSB bloom exceeded 10^9^ cells ml^-1^, among the highest environmental cell densities reported to date, with up to 96 % of the bloom consisting of a single strain-level *Prosthecochloris* lineage (GSB-TRL01). Extreme sulfide concentrations (> 17 mM) coincided with peaking cell density and a shift in isotopic composition. Bloom persistence is supported by tightly coupled sulfur cycling, high rates of nitrogen fixation, mechanisms to tolerate oxidative stress and maintain redox balance, and nutrient acquisition through outer membrane transport systems. A shift in sulfur oxidizing enzymes towards enzymes with higher sulfide affinity proceeded bloom demise. *Prosthecochloris* GSB-TRL01 differs from closely related lineages in several outer membrane transport systems, including porins to transport phosphate and tonB-dependent transporters. Collectively, these findings identify physiological and metabolic strategies that may enable near-clonal *Prosthecochloris* populations to attain extraordinary biomass while coupling the sulfur, carbon, and nitrogen cycles in coastal euxinic environments.

## Introduction

Estuaries and lagoons are highly productive aquatic ecosystems that harbor diverse microbial communities and play an important role in coastal biogeochemical cycling ^1^. Due to the shallow depths of many estuarine and lagoon environments, microbial communities in the sediment and water column are tightly coupled, and input of both fresh and saline water often results in density stratification of the water column ^1^. Stratification, along with high rates of microbial respiration, can result in low-oxygen bottom waters and a shift in microbial community composition and function. As coastal sediment respiration is typically dominated by sulfate reduction ^2^, shallow, oxygen-depleted estuarine and lagoon waters often accumulate sulfide and contain photic, euxinic habitats. Anoxygenic phototrophs, namely purple sulfur bacteria (PSB) and green sulfur bacteria (GSB), are the dominant primary producers in shallow euxinic environments and can form large dense blooms ^3–6^.

GSB (class Chlorobia) in particular are known to form near-clonal blooms that have an important yet understudied role in aquatic carbon cycling. GSB can account for over 80 % of productivity in some aquatic environments ^7,8^ and have outsized roles in sulfide oxidation relative to their population size ^9^. GSB have a high tolerance and affinity for sulfide ^10,11^ and couple sulfide oxidation with carbon fixation using the reverse tricarboxylic acid cycle (rTCA), transforming highly toxic sulfide into various low- to non-toxic reduced forms (e.g. S*^n^_2-_*, S_0_, S_2_O_32-_, SO_42-_). Chlorobi use specialized bacteriochlorophyll pigments (BChl*a*, *c*, *d*, *e*) absorbing far-red light, whose synthesis is inhibited by the presence of molecular oxygen, and unique light-capturing organelles, known as chlorosomes, to capture light energy at extremely low light intensities. Given these capabilities, GSB are typically found in the lowermost photic zone and below the oxygen-sulfide boundary ^10^.

Despite their classification as strict anaerobes ^11^, recent studies have shown that GSB may be able to tolerate some amount of dissolved oxygen (∼ 30 *µ*M), suggesting that their niche may extend into hypoxic waters^5^. GSB can fix nitrogen at high rates ^12,13^, increasing the amount of bioavailable nitrogen and thereby linking the nitrogen, carbon, and sulfur biogeochemical cycles. With increasing oceanic and coastal deoxygenation ^14,15^, potential habitats for GSB are spatially and temporally expanding. Despite the apparent ecological importance of GSB in oxygen-depleted, photic environments, little is known about the mechanisms underlying GSB bloom initiation, the maintenance of dense near-clonal populations over weeks to months, or how these blooms reshape estuarine redox chemistry and biogeochemical cycling.

Seasonal, frequent, and long-lived blooms of GSB occur at shallow depths in Trunk River Lagoon (TRL) estuary of Falmouth, MA (Fig. 1a). The lagoon is brackish with limited turnover, as its southern end is connected to the ocean via a small channel with restricted water flow and the northern end is connected to a kettle pond (Oyster Pond). Bhatnagar et al. studied dense (10^8^ cells ml^−1^) blooms in shallow parts of TRL and showed they were dominated by a single species-level GSB lineage ^5^. GSB blooms form at depths of 10 to 80 cm and up to 30 cm in thickness in deeper parts of the lagoon. Anoxygenic phototrophic blooms at such shallow depths are very unusual, and most previous studies report blooms at depths from 2 to 20 m ^3,6,8,16,17^. Trunk River blooms are thus visible by satellite, and they persist for weeks to months (Fig. 1b). Such naturally occurring blooms of anoxygenic phototrophs are ideal model systems for understanding the physiology of environmental GSB populations, the mechanisms that support microbial bloom formation and maintenance, as well as the effect of estuarine euxinia on microbial community structure and function.

**Figure 1.**
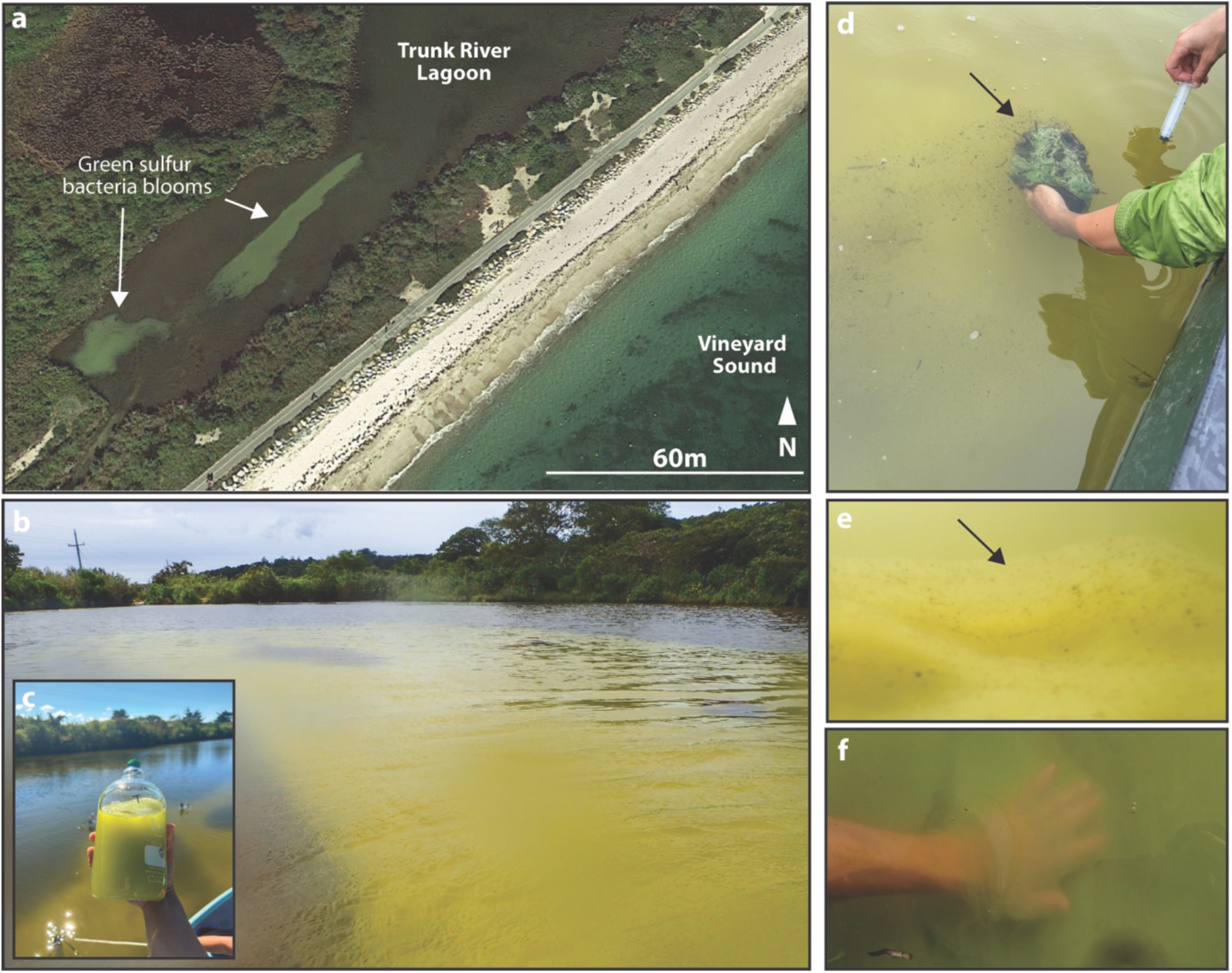
(a) Satellite image of Trunk River Lagoon estuary (Woods Hole, MA) from 2018 (Google Earth) showing occurrence of Green Sulfur Bacteria (GSB) blooms in the southwestern area of the lagoon. (b) Trunk River lagoon during 2021 GSB bloom. (c) Bloom sample collected in a flushed, evacuated anaerobic media bottle. (d) Microbial mat present at the bloom sediment interface. (e, f) Suspended biofilm-like matrix present at interface of bloom with the overlying water column.

To understand the biogeochemical and metabolic dynamics of estuarine GSB blooms, we performed a time series sampling in Trunk River Lagoon during a 58-day bloom period, including 7 consecutive days of sampling during the bloom’s onset and exponential growth phase. Samples were collected from within the bloom and the overlying water column, as well as a microbial mat that formed at the bloom-sediment interface and a suspended biofilm-like matrix that formed at the bloom’s upper interface (Fig. 1). We applied a multi-omics approach (short-read and full-length 16S rRNA metabarcoding, metagenomics, metaproteomics) to understand the diversity and physiology of the bloom from its onset to its demise. We present a comprehensive characterization of the extremely high cell densities, biomass, and activity achieved by the GSB bloom, including stable isotope analysis and water column chemistry, total cell counts, taxa-specific cell counts (CARD-FISH), and both DNA- and protein-based metrics of relative abundance and biomass. Such an extensive methodological approach in environmental microbiology is rarely applied, particularly in a bloom context. This approach offers unique insights into bloom formation and persistence from microbial and biogeochemical perspectives, as well as the rare opportunity to compare interpretations from multiple ‘omics methods and direct cell counts.

## Results and Discussion

### Bloom Dynamics & Biogeochemistry

#### Green sulfur bacteria blooms displayed extremely high cell densities and sulfide concentrations

Cell densities within GSB blooms increased by two to three orders of magnitude over just four days, from 5.3 × 10^7^ ± 9.9 × 10^6^ cells ml^−1^ (Day 2) to 1.2 × 10^9^ ± 1.5 × 10^8^ total cells ml^−1^ (Day 6; Fig. 2a). GSB-specific cell counts from this same period show a rapid increase and subsequent dominance of GSB within blooms, accounting for up to 95 % of total cells (1.1 × 10^9^ ± 1.6 × 10^8^) at peak cell density (Day 6). Our values are to the best of our knowledge the highest total and GSB-specific cell densities ever reported in green sulfur bacteria blooms, which typically contain 10^6^ - 10^7^ cells ml^−1^ and up to 10^8^ GSB cells ml^−1^ ^3,4,12,18,19^. Bacterial cell densities of 10^9^ cells ml^-1^ in a liquid culture or sample are observed primarily in laboratory cultures of fast-growing species, such as *E. coli* ^20^, or in hydrothermal vent waters (10^5^ - 10^9^ cells ml^-1^) ^21^, to our knowledge such densities do not typically occur in coastal environments. The increase in GSB bloom cell density also coincided with a rapid increase in sulfide concentrations from 3.1 mM to 17.5 mM (Fig. 2a), exceeding the highest reported concentrations of sulfide in GSB blooms, which typically range from 100 *µ*M – 5 mM ^3,5,8,18,22–24^. Sulfate concentrations within the bloom ranged from 3.7 mM - 11.9 mM, with sulfide concentrations exceeding sulfate concentrations by nearly 3-fold on Day 6 (6.0 mM sulfate and 17.5 mM sulfide) and nearly 4-fold on Day 7 (3.7 mM sulfate and 14.1 mM sulfide) (Fig. S1). The overall increase in total dissolved sulfur observed is likely attributed to release of sulfide from the sediment to the water column, as well as sulfide production within the bloom from various forms of sulfur (e.g. sulfate, elemental sulfur, polysulfides) (described below).

**Figure 2.**
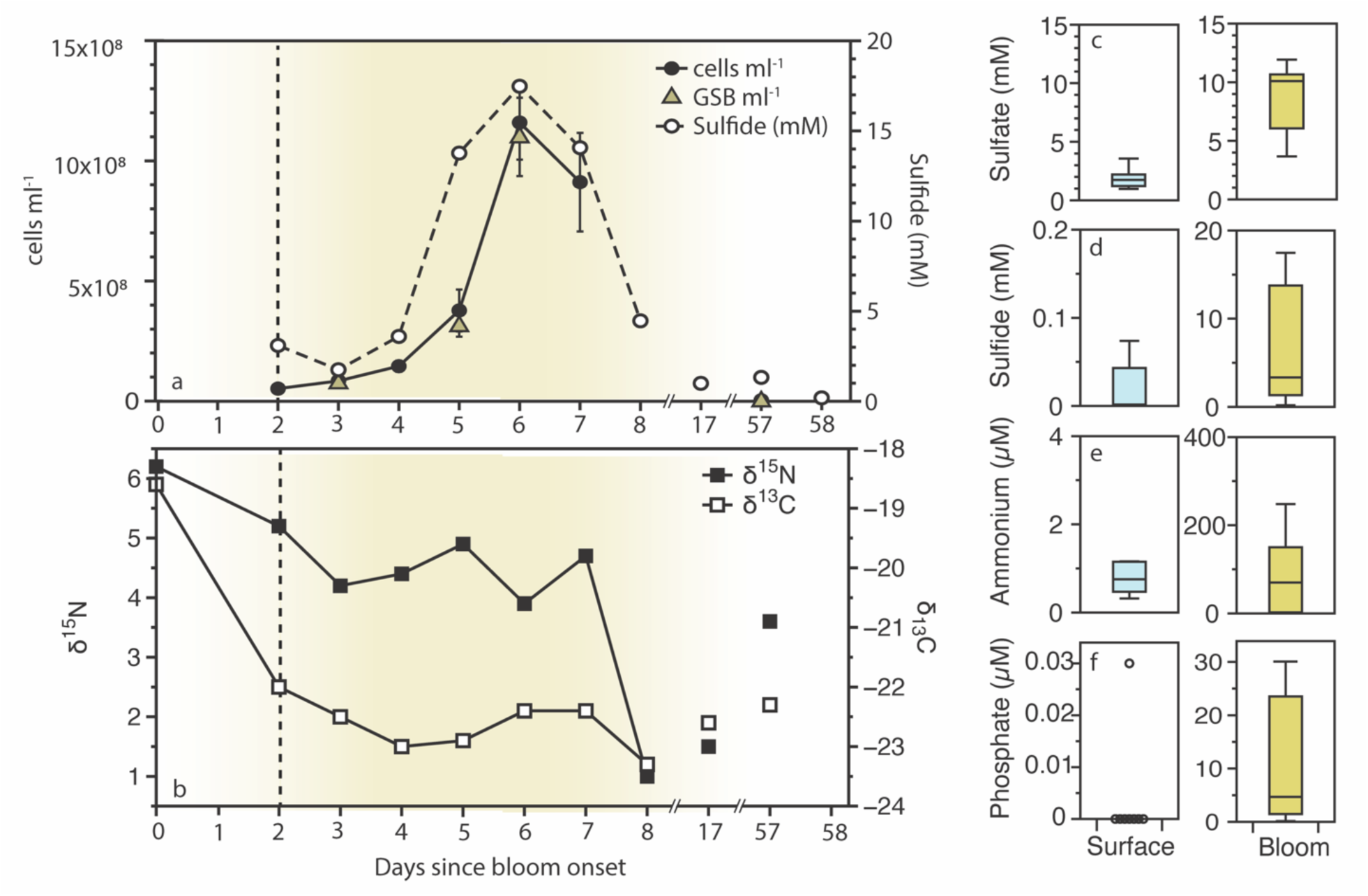
(a) Total cell counts, Green Sulfur Bacteria (GSB) cell counts, and sulfide concentrations (mM) and (b) carbon (δ^13^C) and nitrogen (δ^15^N) stable isotope values of biomass over the duration of a single GSB bloom from August-October 2021 in Trunk River Lagoon estuary. Shading represents the peak bloom period. (c-f) Dissolved inorganic nutrient concentrations of sulfate (mM), sulfide (mM), ammonium (μM), and phosphate (μM) over the duration of the bloom (Day 0, 24-Aug-2021 to Day 58, 21-Oct-2021) from surface water above the bloom and from bloom samples (∼0.4 m depth). Note that y-axes differ on plots d-f with compound concentrations being up to 1000 times higher in the bloom than in the overlying waters. Additional inorganic nutrient concentrations, along with pH, dissolved oxygen, salinity and temperature are shown in supplemental Figs. S2-S4.

#### Nutrient concentrations within and above blooms differ by orders of magnitude

The biogeochemical conditions of the water column differ substantially between the bloom and the overlying non-bloom surface water. Concentrations of sulfide, ammonium, and phosphate are all at least 2 orders of magnitude greater within the bloom than in the surface water (Fig. 2). Sulfate concentrations are approximately 5-fold higher in the bloom than in the surface waters (Fig. 2c). Ammonium is the dominant form of dissolved inorganic nitrogen within the bloom, with nitrate and nitrite concentrations being < 1 μM (Fig. S2 & S3). Ammonium concentrations were highly variable over the bloom period, ranging from 1.9 - 248.0 μM (Fig. S3). Using the ratio of DIN:DIP as a metric of nutrient limitation ^25^ the bloom transitions from a period of nitrogen limitation (DIN:DIP = 4.74, Day 2) to phosphate limitation (DIN:DIP = 54.8, Day 6; Fig. S3). This shift in the state of nutrient limitation coincides with peak cell densities and sulfide concentrations. Elevated sulfide conditions, as well as high organic carbon availability, can favor dissimilatory nitrate reduction to ammonium (DNRA) over denitrification ^26,27^, leading to the production and accumulation of ammonium. Moreover, as nitrogen and carbon fixers, GSB also directly contribute to the production of bioavailable ammonium and organic carbon. A low DIN:DIP ratio (i.e. nitrogen limitation), as observed in the early bloom phase, creates conditions favorable for nitrogen fixation ^28^. Over the development of the bloom, bulk biomass nitrogen isotopic signatures (*δ*^15^N) decreased from 6.2 ‰ to 1.0 ‰ (Fig. 2b) and remained low through Day 17 (1.5 ‰). Low *δ*^15^N values are indicative of N derived from nitrogen fixation, as the *δ*^15^N of atmospheric N_2_ is 0 ‰ ^29^. This decrease in *δ*^15^N is concurrent with a decrease in *δ*^13^C from -18.6 ‰ to -23.3 ‰., with a change of ∼3 ‰ occurring within two days of the bloom onset. GSB have an estimated fractionation factor (ε) of 18 ‰ relative to their inorganic carbon pool ^30^. Measurements of the DIC pool concentration and isotopic composition are necessary to gain a complete picture of the bloom’s effect on carbon fractionation, in order to characterize the carbon source used by GSB for carbon fixation; however, GSB fractionation in an environment with a highly remineralized and isotopically lighter carbon pool could drive the observed decrease in *δ*^13^C. Other community members with larger fractionation factors, such as purple sulfur bacteria (PSB) ^30^, may also contribute to this observed change. These biogeochemical measurements indicate a highly productive and dynamic environment that is sharply distinct from the overlying water column and suggest high rates of both nitrogen and carbon fixation within the bloom.

Water stratification driven by a salinity gradient in the lagoon, as observed in Bhatnagar et al. ^5^ and in this study (Fig. S4), likely helps to maintain the biogeochemical conditions of the bloom. As shown in Bhatnagar et al. ^5^, pH was also lower in the bloom (6.3 ± 0.1, n=6) than in the surface water (6.7 ± 0.4, n=6) (Fig. S4). Lower pH within the bloom is likely caused by the accumulation of dissolved inorganic carbon, driven by high rates of organic matter degradation and sulfate reduction in the lower, more saline waters. Lastly, dissolved oxygen was detected within the bloom, ranging from 4 – 43 µM in the deepest parts of the bloom (Fig. S4), further supporting previous findings of oxygen within GSB blooms ^5^. However, due to the large diameter of the multiparameter probe used here relative to the scale of gradients in ecosystem, these measurements may overestimate the concentration of dissolved oxygen, as dissolved oxygen can be entrained from upper waters during measurement. Microsensors will determine the presence of oxygen within these blooms and its spatiotemporal variability more accurately in further studies.

### Bloom Microbial Composition

#### A single species-level *Prosthecochloris* is extremely abundant in blooms

Blooms were dominated by GSB, which comprised up to 90% - 98% of the population (Day 6; Table 1) across multiple methods of assessing relative abundance. These relative abundances are among the highest ever measured in an anoxygenic phototrophic bloom, where relative abundances of the dominant bloom formers are typically close to 50% ^3,6,31,32^. In the short-read 16S rRNA gene dataset (Illumina, V4-V5), Chlorobiales amplicon sequence variants (ASVs) accounted for up to 96.1% of relative sequence abundance (Day 6) and averaged 82.7 ± 8.2% (n=7) during the development of the bloom (Days 2 - 8; Fig. 3a, Table 1). Relative ASV abundances aligned closely with GSB-specific cells counts from CARD-FISH (Table 1). The lowest alpha diversity was observed on Day 6, coinciding with the maximum cell density (Fig. 2a). Bloom samples were dominated by a single Chlorobiales ASV (ASV1) of the genus *Prosthecochloris*, which represented 94.5 ± 5.8% (n=10) of all Chlorobiales ASVs during the entire bloom period. ASV1 was also 100 % identical to the predominant *Prosthecochloris* variant (SV1) reported in Trunk River Lagoon from sampling in 2015^5^. Within microbial mat and suspended biofilm samples, ASV1 was also the predominant lineage accounting for 84.7 % (mat) and 82.7 % (biofilm) of the community during the exponential phase of the bloom (Fig. S5).

**Figure 3.**
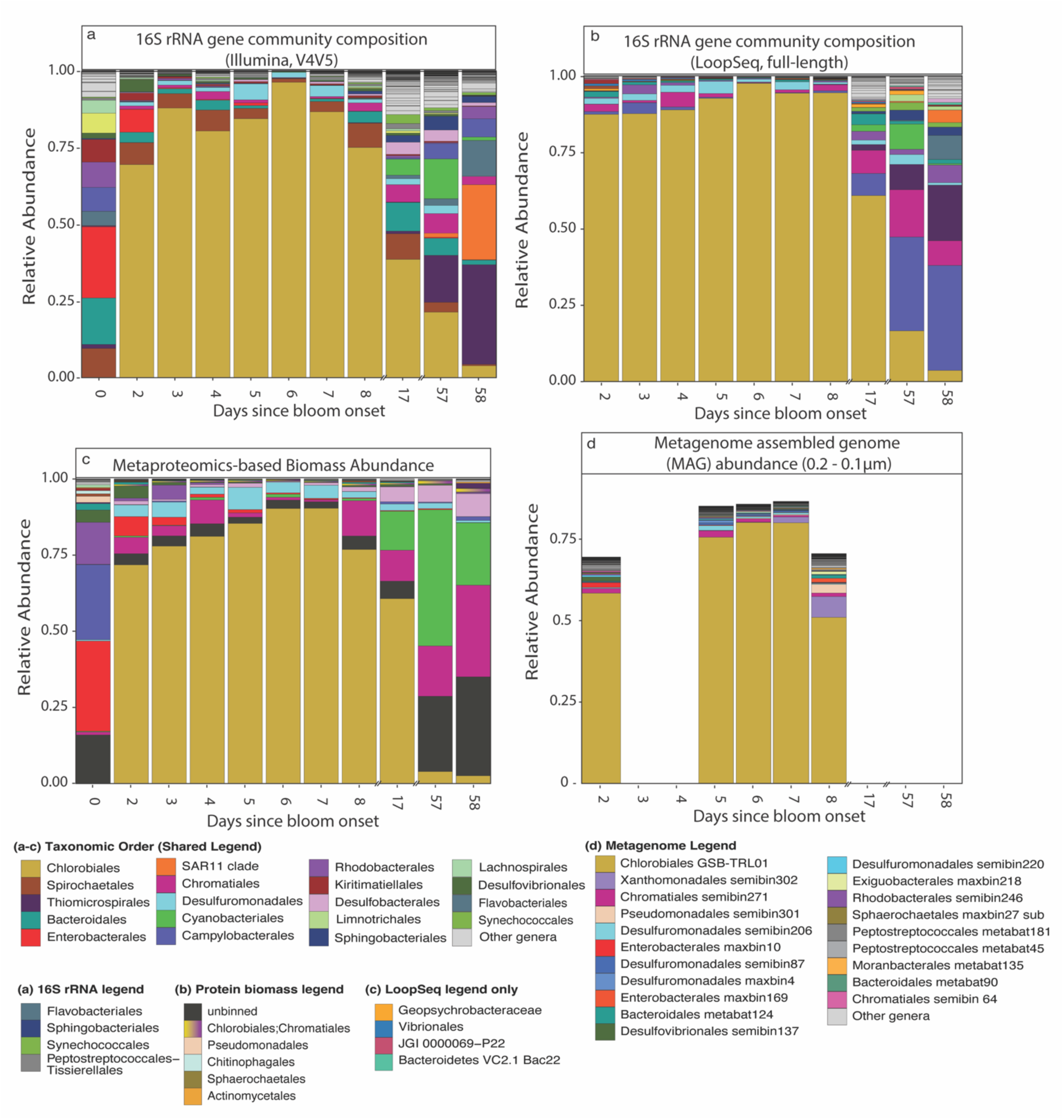
Comparison of abundance measurements by method over GSB bloom period (Day 0, 24-Aug-2021 to Day 58, 21-Oct-2021) with (a) 16S rRNA gene-based relative abundance of the V4-V5 region; (b) LoopSeq full-length 16S rRNA gene-based relative abundance; (c) metaproteomics biomass-based abundance; and (d) metagenome assembled genome (MAG) abundances from samples collected in the 0.2 - 0.1 μm filter size fraction. In plots (a-c), the 20 most abundant taxonomic orders are represented with a shared legend for orders found in multiple plots, and in plot (d) the 20 most abundant MAGs are represented. Note that in for the Protein Biomass Legend, “Chlorobiales;Chromatiales” represents proteins that cannot be distinguished between the two taxonomic groups.

**Table 1.** Cell counts of total cells and GSB-specific cell counts (CARD-FISH) compared with ‘omics-based metrics for measuring community composition: 16S rRNA gene sequencing (V4V5), LoopSeq full-length 16S rRNA gene sequencing, Metagenome-assembled genome (MAG)-based abundance, and metaproteomic-based biomass abundance. Relative abundances of the highly enriched amplicon sequence variant (ASV1) and LoopSeq amplicon sequence variant (L-ASV1) are compared between traditional short-read V4V5 and full-length LoopSeq Illumina methods.

| Days since bloom onset | Date | Total DAPI Cell Counts (cells/ml ± stdev) | CARD-FISH GSB-532 % | Amplicon relative abundance Chlorobiales (%) | LoopSeq relative abundance Chlorobiales (%) | Metagenome abundance (0.1-0.2μm) Chlorobiales (%) | Proteomics Chlorobiales Biomass (%) | Amplicon vs. LoopSeq ASV1 relative abundance (%) | |
| --- | --- | --- | --- | --- | --- | --- | --- | --- | --- |
|  |  |  |  |  |  |  |  | ASV1 | L-ASV1 |
| 2 | 26-Aug-21 | $5.3 \times 10^7 \pm 9.8 \times 10^6$ | - | 69.4% | 87.8% | 58.4% | 71.8% | 68.6% | 70.5% |
| 3 | 27-Aug-21 | $8.5 \times 10^7 \pm 1.4 \times 10^7$ | 90% ± 16.3% | 87.7% | 87.8% | - | 78.0% | 87.3% | 64.9% |
| 4 | 28-Aug-21 | $1.5 \times 10^8 \pm 2.3 \times 10^7$ | - | 80.2% | 89.2% | - | 81.2% | 79.9% | 70.4% |
| 5 | 29-Aug-21 | $3.8 \times 10^8 \pm 8.7 \times 10^7$ | 83.2% ± 12.0 % | 84.3% | 93.0% | 75.5% | 85.4% | 75.2% | 73.2% |
| 6 | 30-Aug-21 | $1.2 \times 10^9 \pm 1.5 \times 10^8$ | 95.5% ± 14.1% | 96.1% | 97.8% | 80.2% | 90.3% | 95.5% | 76.8% |
| 7 | 31-Aug-21 | $9.1 \times 10^8 \pm 2.0 \times 10^8$ | - | 86.5% | 94.6% | 80.1% | 90.4% | 76.3% | 74.3% |
| 8 | 1-Sept-21 | - | - | 74.9% | 95.2% | 51.0% | 76.9% | 74.4% | 74.7% |
| 57 | 20-Oct-21 | $4.1 \times 10^6 \pm 1.6 \times 10^6$ | 21.5% ± 19.5% | 21.3% | 17.3% | - | 4.0% | 18.8% | 11.5% |

In the LoopSeq full-length 16S rRNA gene dataset, Chlorobiales L-ASVs (Loop-ASVs) represented 92.2 ± 4.0% of relative sequence abundance during the initial phase of the bloom (Days 2 - 8), with a single L-ASV (L-ASV1) accounting for 77.2 ± 5.1% of all Chlorobiales L-ASVs in bloom samples during the entire bloom period (n=10; Fig. 3b). This L-ASV1 (1,425 bp) aligned with 100 % consensus against the short-read *Prosthecochloris* ASV1. Alignment of abundant *Prosthecochloris* L-ASV sequences against available full-length reference sequences (MUSCLE 5.1, PPP, in GeneiousPrime)^33^ suggests that *Prosthecochloris* L-ASVs are most closely related to *Prosthecochloris* CB11 (KR013743) with a difference of 4 nucleotides and differ from *Prosthecochloris vibrioformis* (KX417801) by 61 nucleotides (Fig. 4a), representing a difference at the species-level from *P. vibrioformis*.

**Figure 4.**
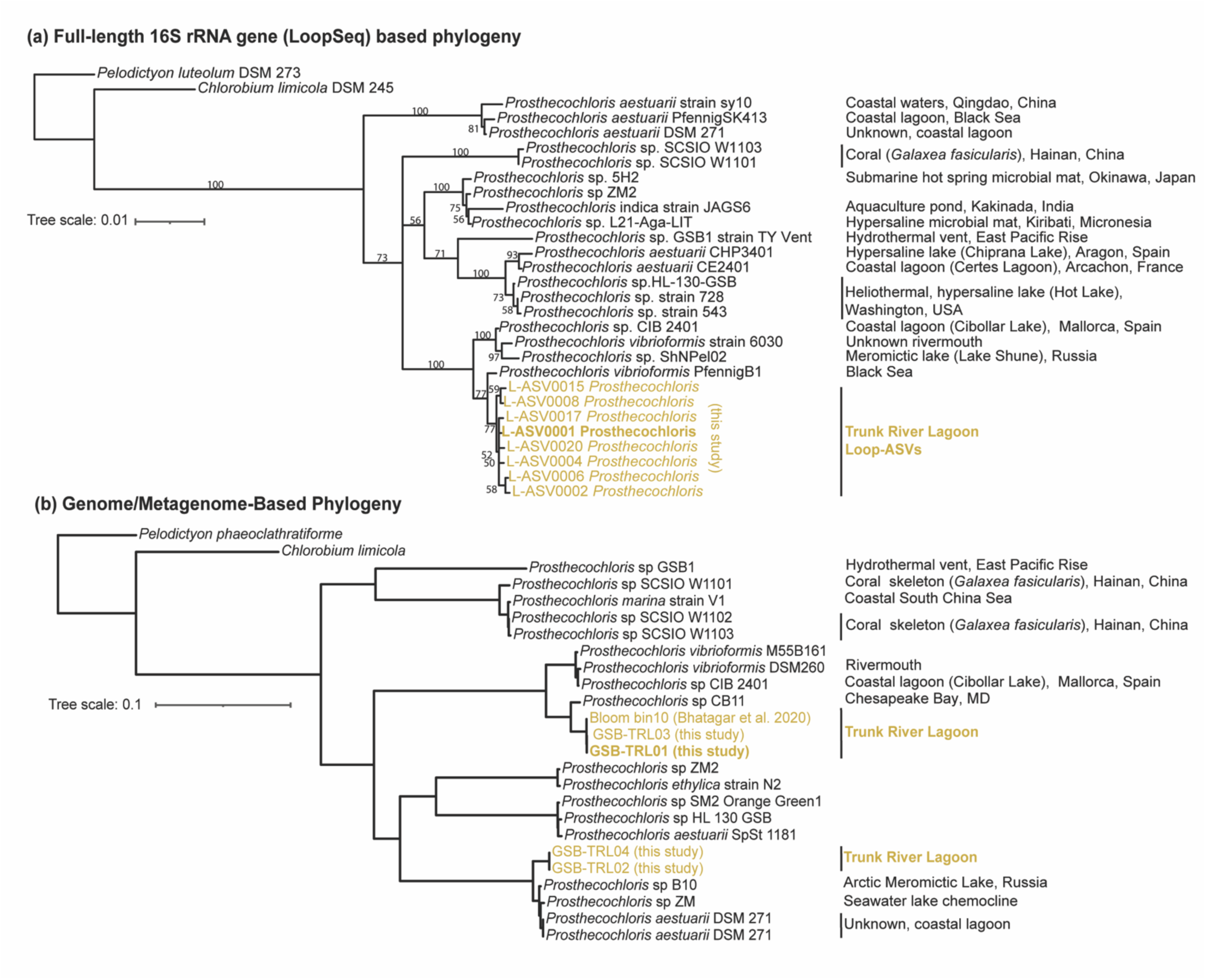
(a) Phylogeny based on LoopSeq full-length 16S rRNA gene sequences of *Prosthecochloris* and those generated in this study using LoopSeq. Sequences were aligned using Multiple Sequence Comparison by Log-Expectation (MUSCLE v5.1) and a maximum-likelihood tree was constructed using Geneious Tree Builder. (b) Genome/metagenome-based phylogeny of *Prosthecochloris* genomes generated using the GTDB-Tk *de novo* workflow including reference sequences and those from this study, as well as the *Prosthecochloris* MAG sampled in 2015 from Trunk River Lagoon ^5^. All accession numbers, fasta files, and tree files are available in the supplementary data repository.

Two *Prosthecochloris* metagenome assembled genomes (MAGs, population genomes) were present in the bloom with one MAG, GSB-TRL01, having the highest coverage (770.3 ± 376.5, n=20) and relative abundance (54.0 % ± 21.5%, n=20) across all samples and time points (Table 2; Figs. 3d & S6). GSB-TRL01 is estimated to be 98.6 % complete with 0 % contamination after manual refinement (anvi-refine), with a length of 2.36 Mb, a GC content of 51.4 %, with 2,266 genes (Table 2). GSB-TRL01 coverage and relative abundance were greatest in the [2 – 0.2 µm] and [0.2 – 0.1 µm] size fractions, with averages exceeding 900 and 68 %, respectively (Fig. S6). The other *Prosthecochloris* MAG (GSB-TRL02) had an average coverage of 0.97 ± 1.0 and average abundance less than 0.2% (n=20) and is estimated to be 97.2 % complete with 2.8 % contamination after manual refinement (anvi-refine), with a length of 2.29 Mb, a GC content of 50.1 %, with 2,209 genes (Table 2). Based on a compilation of 24 *Prosthecochloris* genomes ^34^, GSB-TRL01 and GSB-TRL02 are similar in their metrics in comparison with others of the same genus, where the average genome size, GC content and CDS were 2.4 ± 0.3 Mb, 50.8 ± 3.1%, and 2,217 ± 322, respectively.

**Table 2.** Details of *Prosthecochloris* metagenome assembled genomes (MAGs) from a co-assembly of bloom samples using all size fractions (n=20).

| MAG | Taxonomy | Rel. Abundance | Coverage | Completeness | Contamination | Length | GC Content | CDS |
| --- | --- | --- | --- | --- | --- | --- | --- | --- |
| GSB-TRL01 | *Prosthecochloris* | 54.0 % ± 21.5%, | 770.3 ± 376.5 | 98.6 % | 0 % | 2.36 Mb | 51.4 % | 2,266 |
| GSB-TRL02 | *Prosthecochloris* | < 0.2% | 0.97 ± 1.0 | 97.2 % | 2.8 % | 2.29 Mb | 50.1 % | 2,209 |

GSB-TRL01 appears to be a novel species-level *Prosthecochloris* lineage, with an average nucleotide identity (ANI) of 85.84 with *Prosthecochloris vibrioformis* (GCF_006265245) and falls in the same novel lineage as the MAG ‘Bin10’, which was assembled from the same environment in 2015 ^5^ (Fig. 4b). While *Prosthecochloris* MAGs did not contain 16S rRNA genes, as short-read sequencing struggles to resolve this highly conserved and repetitive region, both phylogenies based on full-length LoopSeq analysis of the 16S rRNA gene and the GSB-TRL01 MAG reveal that *Prosthecochloris* CB11, isolated from an estuarine environment (Chesapeake Bay) ^9,35^ and *Prosthecochloris* CIB 2401, isolated from a coastal brackish lagoon, are closely related lineages (Fig. 4) ^36^. *Prosthecochloris* MAGs from microbial mat samples (from the bloom-sediment interface) reveal that the predominant lineages present in the mat are the same as in the bloom, indicating that there was not a separate benthic population in these GSB mats (GSB-TRL03 & GSB-TRL04, Fig. 4b).

Using a protein-based abundance approach^37^, *Prosthecochloris* GSB-TRL01 comprised 71.8 % of proteinaceous biomass over the entire sampling period, with GSB-TRL02 accounting for only 3.7 % of proteinaceous biomass. Overall, Chlorobiales proteinaceous biomass abundances ranged from 72 - 90 % peak bloom (Fig. 3c, Table 1). DNA-based, protein-based, and direct cell count-based estimates of the relative abundance of Chlorobiales using these different approaches were extremely well-aligned (Table 1), except for the final day of sampling. On Day 58, the decline in the GSB population is more pronounced in the proteinaceous biomass than in DNA-based estimates (Table 1). This likely is in response to more rapid degradation of proteins than DNA in the later phase of the bloom. The ratio of DNA to protein is known to increase due to protein breakdown when cells become nutrient limited and enter stationary phase^38^. Such a comparison of different abundance metrics is rarely performed, as few studies have integrated cell-based, amplicon-based, metagenomic, and metaproteomic abundance metrics for the same environmental population. These results illustrate how each method captures changing growth states and community shifts in a bloom context, and highlight the advantage of proteinaceous biomass metrics in capturing rapidly changing community dynamics. However, while our abundance results are well aligned for dominant taxa during the main bloom period, other minor taxa vary substantially in abundance depending on the method.

#### Abundance of non-*Prosthecochloris* bacteria community members differs with sampling method, highlighting potential biases for minor community members

Among non-Chlorobiales ASVs in the short-read bacterial bloom dataset, the highest relative abundances were found the taxonomic groups Spirochaetales (4.9%), Bacteroidia (2.8%), Desulfuromonadales (2.2%), and Chromatiales (PSB) (1.3%) (Fig. 3a). However, in the LoopSeq full-length (L-ASV) community composition, Spirochaetales (0.3%) and Bacteroidales (0.9%) abundances were lower, while Chromatiales (4.2%) and Thiomicrospirales (3.0%) abundances were greater (Fig 3b). Metaproteomics data reveals high biomass of Chromatiales (8.8%), Cyanobacteriales (8.1%), Desulfobacterales (2.5%), and Desulfurmonadales (2.5%), while Spirochaetales (0.05%) biomass was low (Fig. 3c). In the metagenome dataset, MAGs of the orders Chromatiales and Desulfuromonadales were abundant (Fig. 3d). All methods clearly suggest that Chromatiales and Desulfuromonadales are important minor clades within the bloom. However, differences among other populations could lead to varying interpretations of non-Chlorobiales taxa. Notably, based on short-read amplicon data alone, Spirochaetales appear to be a major constituent of the bloom, yet this is not supported by full-length LoopSeq or metaproteomics data. This could be a result of primer bias in the V4V5 primer pair rather than relic DNA, as relic DNA would presumably have also been captured in the LoopSeq and metagenome datasets.

Additionally, both Cyanobacteria and Chromatiales clades, which contain sulfur-oxidizing and carbon and nitrogen fixing taxa (discussed below), contribute substantially more to the bloom protein biomass than DNA-based sequencing metrics. Cyanobacteria have been shown to have a higher contribution to metaproteomes compared with amplicon sequencing abundance methods ^37^, which may be due to their larger size. In the Trunk River GSB bloom, filamentous *Geitlerinema* were the most abundant cyanobacterial taxon, which can have volumes up to 33 µm^3 39^. Cells of this size would be approximately 2 orders of magnitude larger in volume than *Prosthecochloris*, with an estimated volume of 0.2 µm^3^. Similarly, abundant Chromatiales taxa, such as *Marichromatium*, can have cell volumes of ∼7 µm^3^ ^40^, an order of magnitude larger than *Prosthecochloris*. The differing abundances of these two clades in the metaproteome compared with DNA-based methods illustrates the differing perspectives that these techniques provide on a community. While DNA-based methods provide insight into cell counts, they do not accurately reflect biomass or potential activity. In the case of Trunk River, these findings suggest that both Cyanobacteria and Chromatiales are active within the community, have large cell sizes relative to their population, and play an important role in the bloom and its biogeochemistry.

In other blooming systems, heterotrophic bacteria increase in abundance during bloom demise, suggesting that they remineralize carbon fixed by the bloom ^41^. Although our sampling period was not optimized to examine the question of a secondary heterotrophic bloom developing during the decline of the GSB bloom, we do see an increase in absolute cell numbers of non-GSB cells, as calculated by subtracting GSB specific counts from total cell counts during the bloom (from 8.5 x 10^6^ cells mL^-1^ three days after bloom onset to 6 x 10^7^ cells mL^-1^ six days after bloom onset). After the bloom, non-GSB cell counts decline to 3.3 x 10^6^ cells mL^-1^, suggesting that the bloom fuels a larger population of heterotrophic bacteria, which likely contribute to the remineralization of fixed carbon.

#### Blooms may support populations of acidophilic and methanogenic archaea

In addition to non-*Prosthecochloris* bacterial community members, we also report the archaeal community composition of the bloom using archaeal specific primers, marking the first known documentation of potential GSB bloom-associated Archaea. Acidophilic Thermoplasmata (Marine Benthic Group D and DHVEG-1) and methanogens (Methanomicrobia & Methanosarcinia) were both abundant, as well as Nanoarchaeota (Woesearchaeales). Aenigmarchaeota and Asgardarchaeota were also present at > 3 % of the archaeal community during the main bloom period. Between early and late bloom periods, relative sequence abundance of Thermoplasmata and Methanomicrobia increased from approximately 30 % to 45 % and 16 % to 26 % respectively of the archaeal community, while the relative sequence abundance of Methanosarcinia decreased from 16 % to 4% (Fig. S7). As Methanomicrobia are known hydrogenotrophic methanogens, they could increase at the end of the bloom period in response to organic matter degradation and H_2_ production. However, only one archaeal MAG was recovered from the metagenomic dataset, identified as Woesearchaeales UBA583 (Nanoarchaeota, 88% complete, 2.7% contamination, 1.3 Gb) and no Woesearchaeales were present in the metaproteomics biomass bloom data. It is possible that archaeal proteins were present in the unbinned fraction of the dataset, or that archaeal DNA within the bloom is originating from the sediment, where conditions may be favorable for methanogenesis. However, given the relatively low sulfate concentrations during peak bloom days (<5 mM), and the metabolic versatility of some methanogens, it is possible that the bloom itself could be a favorable environment for methanogenesis. Measurements taken in 2024 suggest that these GSB blooms could contain up to 68 µM of methane (unpublished data). Given the importance of methane emissions in coastal environments, future studies should investigate the role of methanogens in these shallow blooms and the underlying sediment.

### Prosthecochloris bloom physiology

#### *Prosthecochloris* express multiple sulfur oxidizing enzymes

As sulfur-oxidizing phototrophs, GSB depend on a source of reduced sulfur to oxidize. Proteomics data of S-cycling enzymes throughout the bloom timeseries shows that indeed oxidative enzymes from GSB, sulfide quinone reductase (Sqr) and sulfide dehydrogenase (FccB), which oxidize sulfide to polysulfide/elemental sulfur, were abundant during the peak bloom period (Fig. 5). *Prosthecochloris* in the bloom contain Type IV Sqr (SqrX) (Fig. S8), which is commonly found in GSB^42^. While some Type IV Sqr enzymes have high sulfide affinity, enabling GSB to rapidly uptake sulfide even at low concentrations, others are adapted to high sulfide concentrations ^43^, as is likely the case with the SqrX found in GSB-TRL01, a population that thrives at extremely high sulfide concentrations (>2 mM, Fig. 2). In addition to Sqr and Fcc, polysulfide reductase/thiosulfate reductase (PsrA/PhsA) from *Prosthecochloris* was also abundant, accounting for ∼25 % of total S-cycling proteins (Fig. 5), as were the subunits PsrB/PhsB and PsrC/PhsC (Fig. 6).

**Figure 5.**
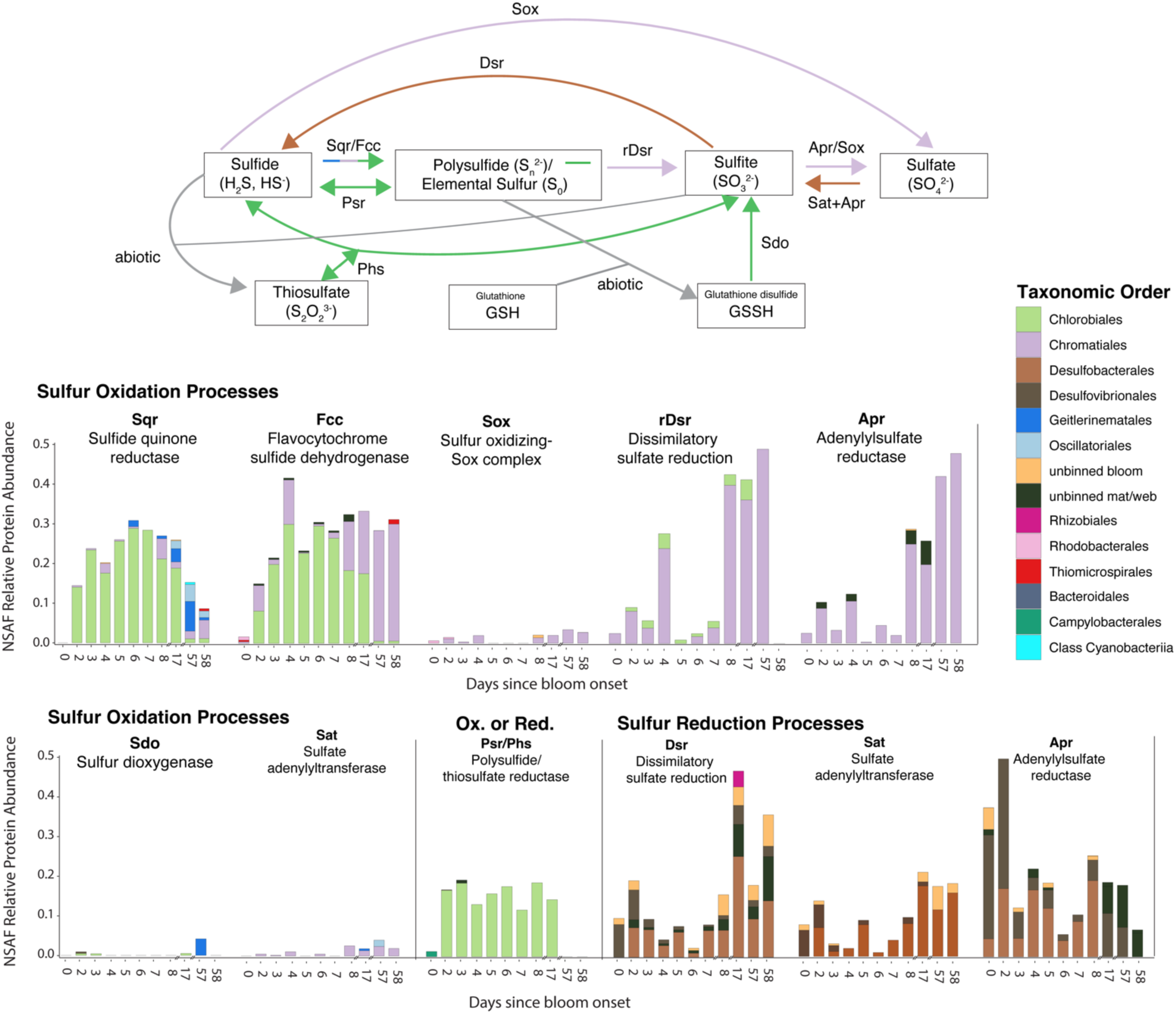
Normalized spectral abundance factor (NSAF) protein abundance of sulfur cycling proteins found in the entire Trunk River GSB bloom community across the timeseries (26-Aug-2021 to 21-Oct-2021) and bloom samples (i.e. water column). Proteins are organized by sulfur oxidation processes and sulfur reduction processes.

**Figure 6.**
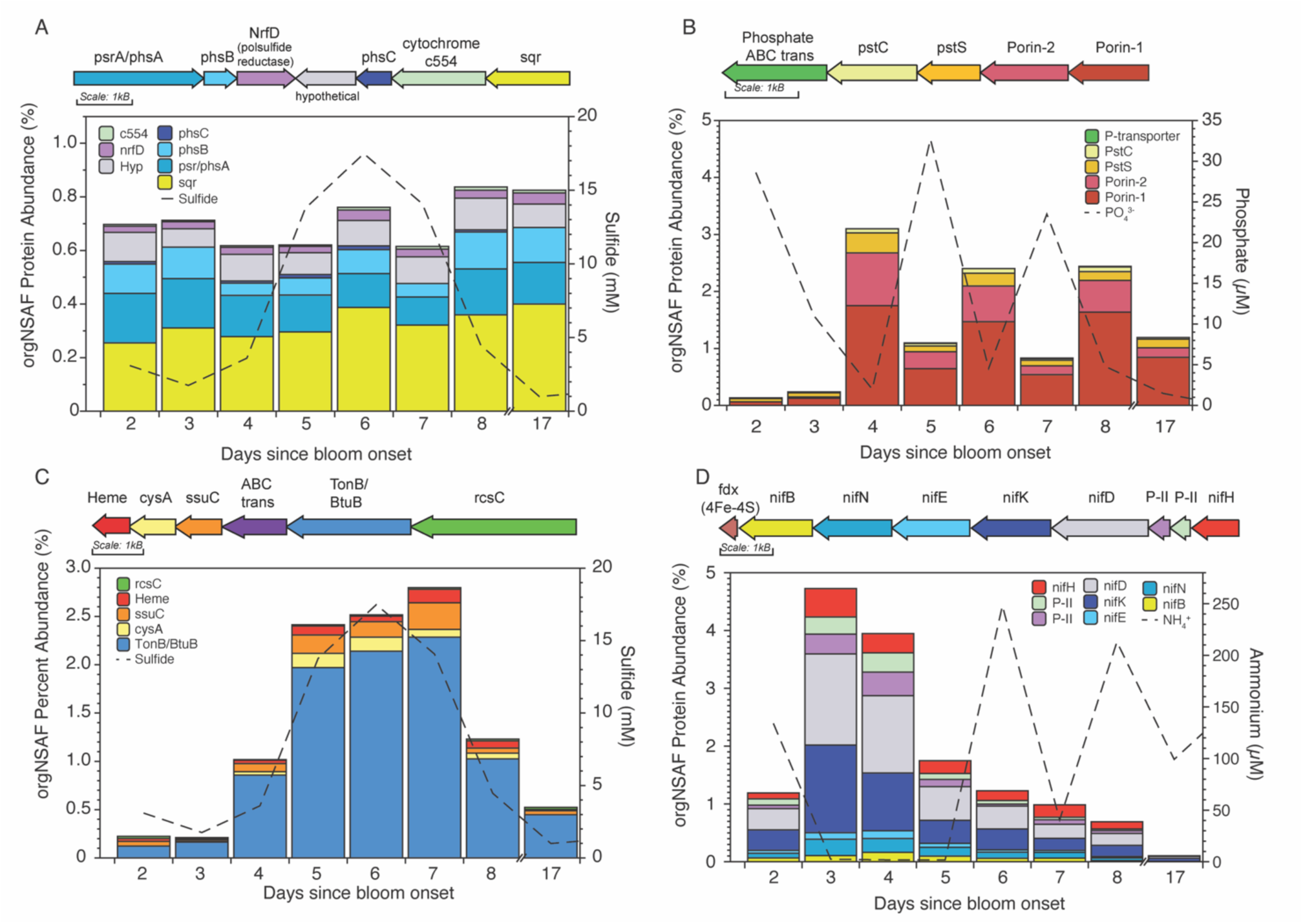
Organism normalized spectral abundance factor (GSB-TRL01*-*NSAF) abundance of gene clusters of interest from *Prosthecochloris* GSB-TRL01 over the main period of the bloom (days 2-16). Gene clusters contain (a) sulfur cycling genes sulfide quinone reductase (sqr) and polysulfide/thiosulfate reductase (psr/phs); (b) phosphate transporters and putative phosphate-related porins; (c) a highly abundant tonB/btuB type outer membrane transporter; and (d) a fully functional nitrogenase. Panels ABCD in this figure correspond to regions outlined in red in Figure 7.

Polysulfide and thiosulfate reductases are complex iron–sulfur molybdoenzymes with few characterized representatives whose catalytic subunits (PsrA/PhsA) share a close phylogenetic relationship and are difficult to distinguish based on their sequences ^44,45^. Polysulfide-reductase-like complexes have been found in numerous GSB genomes ^46^. In *Chlorobaculum tepidum*, a putative polysulfide oxidoreductase (PsrABC) was upregulated in response to sulfide addition, and it has been hypothesized to play a role in either extracellular S(0) globule formation—by oxidizing polysulfide to S(0)—or in S(0) consumption through reduction ^35,47^. In the case of the oxidative direction, Psr would oxidize polysulfide, depositing S(0) and shifting the sulfide pool towards more reduced species. In the reductive hypothesis, sulfide generated by the reduction of polysulfide would accelerate the consumption of extracellular S(0), theoretically generating even more sulfide and facilitating access to S(0) in cells without direct contact to this S pool ^35,48^. These opposing hypotheses—oxidative or reductive—illustrate the uncertainty surrounding the function of Psr/Phs enzymes in GSB. In *Prosthecochloris* CB11 and CIB 2401, a polysulfide-reductase like complex was present in the same gene cluster as Sqr, similar to what is observed here in GSB-TRL01 ^35^. Despite the limits of distinguishing the substrate and function of PsrA/PhsA phylogenetically, GSB-TRL01, CB11 and CIB 2401 all have a closely related PsrA/PhsA enzyme within the same lineage, which differs phylogenetically from the Psr of *C. tepidum* and other GSB species (Fig. S9). While its function in GSB-TRL01 is unknown, given the proximity of this putative polysulfide reductase to the Sqr gene as well as its abundance in the organism normalized proteome (Fig. 6), it may play a role in further oxidation of sulfur compounds within the bloom and the accumulation of S(0). However, it is also possible it plays a reductive role, given that sulfide concentrations increase concomitantly with the GSB population, yet other reductive sulfur enzymes from Desulfobacterales and Desulfovibrionales were not abundant until the later stages of the bloom (Fig. 5). It has previously been observed that a large fraction of sulfide consumed within anoxygenic phototrophic blooms is produced within the bloom itself ^17^, and that sulfate reducing bacteria alone are unlikely to be able to provide the requisite sulfide to fuel a bloom^32^.

#### A shift in S-cycling enzyme abundances precedes community shift and bloom termination

On Day 8 after the bloom onset, the contribution of PSB (Chromatiales) to oxidative S-cycling proteins increases 14-fold through the combination of flavocytochrome sulfide dehydrogenase (Fcc), reverse dissimilatory sulfate reductase (rDsr), and adenylyl sulfate reductase, which collectively would transform sulfide to sulfate with polysulfides/elemental sulfur and sulfite as intermediates (Fig. 5); the abundance of cyanobacterial sulfide quinone reductase (Sqr) also increases (Fig. 5). The relative increase in both PSB and cyanobacterial S-oxidizing proteins precedes the increase in PSB and cyanobacterial DNA-based abundance metrics (Fig. 3) and occurs when GSB cell densities are still high (Fig. 2). Notably, oxidation of sulfide by PSB sulfide dehydrogenase (Fcc) increases at the end of the bloom, while the overall sulfide quinone reductase (Sqr) abundance in the entire community decreases (Fig. 5). Sqr is membrane bound whereas Fcc is located in the periplasm, giving it a higher affinity for sulfide ^49–51^. Sqr types also differ in their sulfide affinities. In Trunk River bloom samples, PSB and cyanobacteria each contained two different Sqr types: Type VI (SqrF) & Type IV (SqrD) in PSB and Type VI (SqrF) & Type I (SqrA) in cyanobacteria (Fig. S8). SqrF and SqrA are known to have high sulfide affinity ^52^, and may be able to outcompete the GSB’s SqrX, which is likely adapted to high sulfide concentrations, when sulfide concentrations decrease in the final stages of the bloom. This suggests that the demise in the GSB bloom could be linked to a shift in sulfur speciation and/or differing enzyme affinities for sulfur substrates among taxa.

Interestingly, the majority of sulfur cycling proteins within the microbial mat during the peak bloom period (Days 6-8) were from Cyanobacteria and PSB (Chromatiales) (Fig. S10). While there was also some contribution of *Prosthecochloris* to sulfur cycling within the mat, it did not correspond with the mat’s community composition during these days (88 % *Prosthecochloris*) (Fig. S5). This suggests that these two phototrophic taxa, which typically occupy vertical niches above GSB, due to oxygen and sulfide requirements ^53^, were more active in sulfur cycling below the dense GSB bloom than above or within the bloom. Limited photons of light likely reach this mat, given the density of the overlying GSB bloom (10^9^ cells ml^-1^), yet these lineages still appear to perform anoxygenic phototrophy and to outcompete *Prosthecochloris* for sulfide at the sediment interface.

#### Outer membrane transporters play a key role in *Prosthecochloris* GSB-TRL01 bloom physiology

In addition to electron donors, *Prosthecochloris* must rapidly uptake essential nutrients from its environment to form and sustain such dense blooms. The proteome of aligned GSB-TRL01 over the course of timeseries sampling provides further insight into proteins that are critical to the success of this organism during blooms, highlighting the importance of transporters (Fig. 7). Porins and tonB-dependent (TBDT) transporters were highly abundant and accounted up to 4 % of the organism normalized proteome (Fig. 7). Both porins and TBDTs are outer membrane proteins that are associated with gram negative bacteria, such as GSB. Porins facilitate the transfer of small hydrophilic nutrients (sugars, amino acids, inorganic ions) ^54,55^ while TBDTs transport larger/scarce nutrients (vitamin B12, iron complexes) into the periplasm ^56^. TBDTs, however, are a diverse family of transporters and can access a variety of substates, including carbohydrates, amino acids, and organic acids ^56–60^. The expression of TBDTs has been linked to microbial blooms and shown to shift throughout bloom phases and substrate availability^59^.

**Figure 7.**
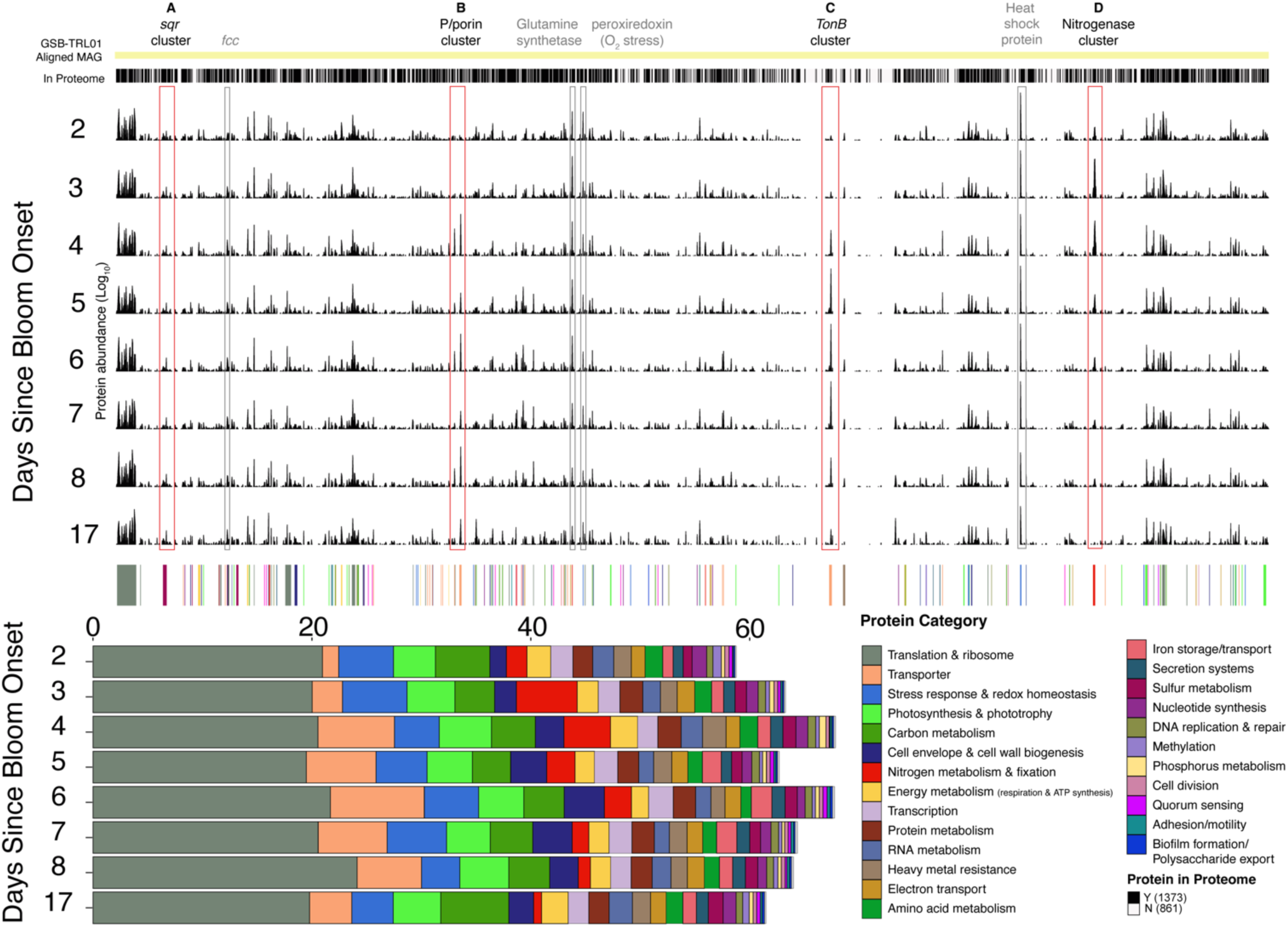
Aligned metagenome-assembled genome (MAG) and proteome of *Prosthecochloris* GSB-TRL01 showing proteins that were found in the proteome and organism normalized spectral abundance factor (GSB-TRL01*-*NSAF) abundance (log10-scale) over time during the primary 17-day sampling period of the bloom. The 250 most abundant proteins were categorized and are plotted using GSB-TRL01 organism normalized NSAF abundance (non-log scale) on the bar plot below. Abundant proteins of interest are outlined. Proteins outlined in red correspond to panels A-D Figure 6, including the sulfide quinone reductase (sqr) gene cluster, the putative phosphate porin gene cluster, the most abundant tonB-dependent (TBDT) transporter gene cluster, and the nitrogenase gene cluster. Proteins not shown in Figure 6 and highlighted in grey include: flavocytochrome sulfide dehydrogenase (fcc), glutamine synthetase, peroxiredoxin, and a heat shock protein (60 kDa chaperonin).

Three TBDTs were abundant in the metaproteome, and based on neighboring genes, at least two appear to be involved with the transport of vitamin B12 in different gene clusters. One of these TBDT-containing gene clusters in GSB-TRL01 encodes a suite of B12 biosynthesis and uptake genes (cobADQPT, cbiDJGFETCHLKp, BtuBDF) (Fig. 8a), while the other is adjacent to multiple cobN gene copies. B12 biosynthesis is a complex process that involves over 20 genes ^61^. CbiG-cbiF and cbiC-cbiH in this region appear to be cbi fusion proteins ^62^, with amino acid lengths and annotations that correspond to multiple protein domains. De novo B12 biosynthesis is not ubiquitous among prokaryotes or GSB, and not all *Prosthecochloris* can synthesize B12, including GSB-TRL01’s closest relative, CB11 ^35^ (Fig. 8a). In GSB-TRL01’s B12 biosynthesis and uptake cluster, the cbi enzymes are part of an anaerobic-type B12 de novo biosynthesis pathway, while the cob and btu enzymes are associated with downstream biosynthesis and/or salvage pathways ^61^. Cbi proteins in this cluster represented approximately 0.2% of the GSB-TRL01 organism-normalized spectral abundance factor (GSB-TRL01-NSAF), while cob and btu proteins in this cluster were ∼0.1% GSB-TRL01-NSAF and ∼0.7% GSB-TRL01-NSAF, respectively. The presence of these proteins indicates that both synthesis and uptake/salvage were occurring during the bloom period. Vitamin B12 is essential for growth and may also be involved in stress response. GSB-TRL01’s proteome contains a Class II vitamin B12-dependent ribonucleotide reductase (0.15 ± 0.01 % GSB-TRL01-NSAF), which may aid in DNA repair during environmental stress. In other GSB blooms, coverage of genes related to B12 biosynthesis and salvage has been observed to be seasonally dynamic, increasing during the summer months when rates of photosynthesis and cell densities are highest ^63^. Our observations confirm that B12 uptake and biosynthesis are essential to GSB-TRL01’s ecological success.

**Figure 8.**
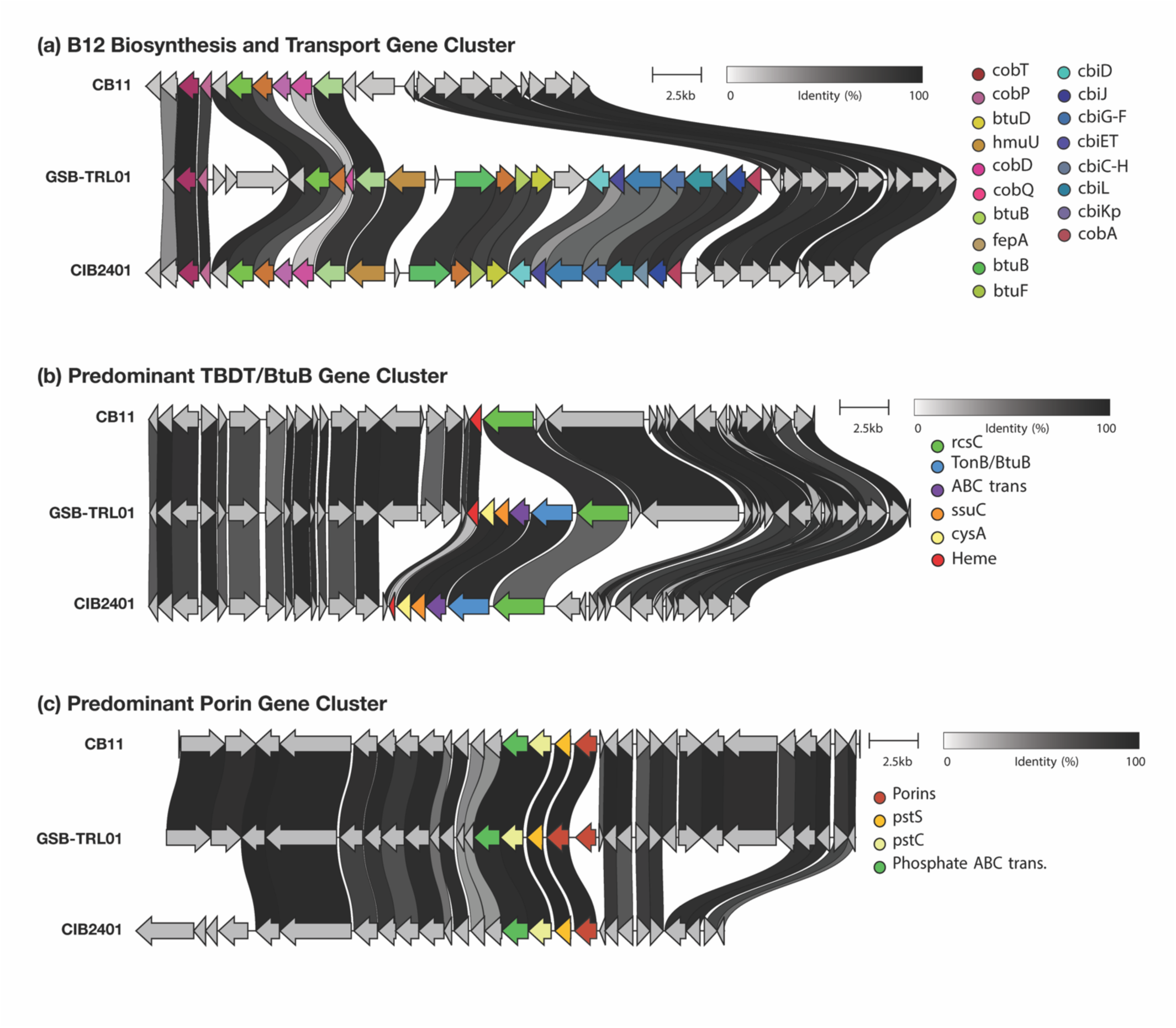
Gene cluster of regions containing (a) vitamin B12 transport and biosynthesis genes, (b) the highly abundant TonB dependent transporter (TBDT) protein and (c) highly abundant porin proteins in the GSB-TRL01 metagenome assembled genome from Trunk River Lagoon. Synteny is shown of closely related *Prosthecochloris* strains, CB11 and CIB 2401. B12 genes are mostly absent in CB11 and present in CIB 2401. The TBDT gene cluster is absent in CB11, and GSB-TRL01 appears to contain and additional phosphate-related porin.

While TBDTs, like BtuB, were previously thought to be restricted to iron and vitamin B12 transport, they are now known to transport a much wider range of substrates (e.g. nickel, cobalt, copper, maltodextrins, sucrose, thiamin and chito-oligosaccharides) ^56^ and play an important role in microbial nutrient acquisition ^64^. The most abundant TBDT in the proteome was not located next to any vitamin B12 related genes, and its abundance increased rapidly from days 3 to 4, reaching up to 2.3 % GSB-TRL01-NSAF (Figs. 6–7). Its organism-normalized spectral abundance factor (GSB-TRL01-NSAF) closely followed the dynamics of both GSB cell density and sulfide concentrations over the course of the bloom, indicating that the number of TBDT transporters per GSB cell increased during the bloom’s exponential phase. Neighboring genes in this gene cluster include ABC transporter proteins involved in organic sulfur transport (cysA, ssuC, sulfonate ABC transporter; Fig 6c). Synteny analysis shows that this gene cluster is absent from closely related CB11 ^35^, while present in more distantly related CIB 2401^36^ (Fig. 8b). Experimental work on GSB *Chlorobium vibrioforme* found that sulfide oxidation was coupled to the reduction of organic sulfur compounds (DMSO). This was hypothesized to protect cells from toxic sulfide concentrations and help dispose of excess electrons ^65^. The authors further suggest that anoxygenic phototrophs may reduce other organic sulfur compounds, in addition to DMSO ^65^. In Trunk River Lagoon, rapid sulfurization of organic compounds has been documented within GSB blooms (D. Dumit, *accepted at Organic Geochemistry*). Given the abundance of this TBDT in the proteome, its neighboring genes involved in inner membrane transport of organic sulfur compounds, as well as the highly sulfidic environment in which this lineage is found, this gene cluster could be involved in the transport of organic sulfur compounds that are subsequently reduced and coupled with sulfide oxidation. This could represent a potential electron sink to maintain cellular redox balance in this highly sulfidic and shallow environment, where high-light conditions and abundant electron donor availability could create an excess of reducing equivalents in the cells.

#### Putative phosphate uptake porins in *Prosthecochloris* respond rapidly to environmental conditions

In addition to these TBDTs, two porins represented over 2 % of GSB-TRL01’s organism normalized proteome. These porins were found within a single gene cluster and located adjacent to phosphate transport systems in the genome (pstC, pstS, phosphate ABC transporter; Fig. 6b). Protein abundance in this gene cluster showed an inverse correlation with bloom phosphate concentrations, suggesting that these porins are involved in phosphate transport and that cells were actively and rapidly modifying the porin content of their outer membranes in response to external phosphate concentrations. Moreover, in comparison with its two closest relatives, CB11 and CIB 2401, the GSB-TRL01 *Prosthecochloris* lineage encodes an additional porin protein (Porin-2) in this same gene cluster (Figure 8c). The ability to rapidly regulate phosphate acquisition in response to environmental conditions may confer an important advantage to this lineage.

#### Prosthecochloris modulates ratio of nitrogenase protein subunits during low ammonium period of bloom

Nitrogenase enzymes were highly abundant within the *Prosthecochloris* GSB-TRL01 proteome (Figs. 6–7), suggesting that access to ammonium is critical for the bloom. Substantial N-fixation is also supported by the natural abundance N-isotope data (Fig. 2). The structural components of the nitrogenase enzyme are encoded by nifDK (Component I), a heterotetrametric complex that contains the active site for N_2_ reduction, and nifH (Component II), a homodimeric dinitrogenase reductase that couples ATP hydrolysis to electron transfer ^66^. Over the course of the bloom, nitrogenase proteins were highly abundant, reaching a maximum 3 days after the bloom onset, and nifDK and nifH from *Prosthecochloris* GSB-TRL01 accounted for 2.6 % and 0.3 %, respectively, of total proteins within the bloom. Peak nitrogenase abundance on days 3-5 of the bloom coincided with lower ammonium concentrations (< 3 *µ*M; Fig. 6d) and a rapid increase in GSB cell density, from 10^7^ to 10^8^ cells ml^−1^ (Fig. 2).

The large increase in nitrogenase abundance during the bloom was driven by an increase in *Prosthecochloris* nifDK, where nifD and nifK each accounted for ∼1.5 % of the *Prosthecochloris* GSB-TRL01 organism-normalized proteome, whereas nifH accounted for only 0.5% (Fig. 6d). This discrepancy in the relative abundances of the nitrogenase structural proteins is unusual, as nifH and nifDK are typically found in similar abundances or with greater nifH than nifDK ^67,68^. The difference between nifH and nifDK abundance could be a result of oxidative stress within the bloom, as the oxygen sensitive Fe-S clusters of nifH are near the protein surface and thus more vulnerable to oxygen than the FeMo cofacter in nifDK ^69–71^. *Prosthecochloris* contains a nifH protein from Cluster III, which is known to be associated with obligate anaerobes ^66,72^ (Fig. S11). Both direct measurements of dissolved oxygen within blooms and abundant peroxiredoxin proteins, accounting for 1 % of the *Prosthecochloris* proteome on Day 2 (Fig. 7), suggest that oxygen stress is occurring in the bloom. Additionally, the high abundance of a 60 kDa chaperonin heat shock protein (3.4 % GSB-TRL01-NSAF; Fig. 7), which assists with protein folding, can also be indicative of oxidative stress ^73–75^. Alternatively, a single nifH in *Prosthecochloris* could associate with multiple nifDK clusters, as nifH is known to transiently bind and dissociate from nifDK when transferring electrons ^76^. This recycling of nifH could reflect a strategy to enhance the efficiency of nitrogen fixation by this lineage.

While direct rates of nitrogen fixation were not measured in this study, high rates of nitrogen fixation within the bloom are supported by nitrogen stable isotope results. The N isotopic signature of biomass shows a complete shift in the bloom community from having a typical coastal environment signal (5.2 ‰, Day 2) to having one derived from atmospheric fixed nitrogen (1.0 ‰, Day 8) (Fig. 2). While many members of the community fix nitrogen, *Prosthecochloris* GSB-TRL01 nitrogenase proteins alone accounted for over 50 % of all nifHDK proteins (Fig. S12). Additionally, Glutamine synthetase (GS), which converts ammonium fixed by nitrogenase into organic nitrogen, was highly abundant (Fig. 7), accounting for up to 1.78 % NSAF of the normalized *Prosthecochloris* GSB-TRL01 proteome (GSB-TRL01-NSAF). By removing freely available ammonium, GS indirectly supports nitrogen fixation by acting as an ammonium sink and preventing an ammonium “shut-off” of nitrogen fixation, which has been documented in GSB ^77–79^. Nitrogen fixation, particularly during the exponential phase of bloom growth, likely allows *Prosthecochloris* GSB-TRL01 to keep up with the nutrient demand that such a dense bloom requires. Nitrogen fixation appears to continue through the bloom period, even when ammonium concentrations are elevated, supporting findings that substantial nitrogen fixation can still occur under nitrogen replete conditions ^80,81^.

## Conclusion

By combining multi-omics, microscopy, stable isotope analyses, and biogeochemical measurements, we comprehensively characterize the metabolisms and physiological adaptations that sustain one of the densest green sulfur bacterial blooms reported to date. We used a unique combination of microbial abundance metrics to gain insights into the community structure at unprecedented resolution and robustness. Our work reveals metabolic strategies that explain the extreme microbial productivity of blooms under euxinic conditions and underscore the capacity of shallow, stratified systems to transform coastal biogeochemistry.

Beyond documenting bloom dynamics, our results offer mechanistic insight into how blooms of *Prosthecochloris* (GSB-TRL01) form and persist. The abundance and co-localization of sulfide oxidation (Sqr) and polysulfide/thiosulfate reductase proteins (Psr/Phs), together with the simultaneous peak of sulfide concentration and cell density, suggests that these enzymes play a key role in the bloom’s success. In the oxidative direction, Psr/Phs would cause an accumulation of elemental sulfur that could fuel further oxidation or reduction. These findings explain previous studies, which suggest that sulfur is produced within blooms and that sulfate reduction can be decoupled from sulfide concentrations.

Additionally, *Prosthecochloris* proteins involved in nutrient uptake and acquisition were extremely abundant and dynamically responded to environmental conditions, including B12 biosynthesis genes, outer membrane TonB-dependent transporters (TBDT), and additional phosphate-associated porin absent in closely related lineages, as well as structural components of the nitrogenase enzyme. Neighboring genes of the most abundant TBDT suggest a role in the transport of organic sulfur compounds. Organic sulfur reduction could be coupled with sulfide oxidation as an additional electron sink to maintain redox balance in such a shallow and highly sulfidic environment. These dynamics highlight the importance of resource acquisition and redox balance during rapid bloom expansion and suggest rapidly scaling up transport capacity is a key component of bloom success. In later phases of the bloom, sulfur cycling protein abundance shifts towards higher affinity sulfide oxidation enzymes of PSB and Cyanobacteria. This shift precedes the DNA-based community change in the relative abundance of PSB and Cyanobacteria, indicating that functional succession occurred before taxonomic succession. We also observe an increase in non-GSB cells at the height of the bloom, suggesting that the bloom supports higher cell numbers of heterotrophic bacteria which act to remineralize fixed carbon. Together, these results demonstrate that multiple metabolic and resource acquisition strategies contribute to the success of *Prosthecochloris* GSB-TRL01 blooms in Trunk River Lagoon, and that comparison with closely related lineages can be used to understand its potential advantages in this environment. As coastal hypoxic and anoxic environments expand globally and microbial blooms increase in frequency, understanding the physiological mechanisms involved in bloom formation, persistence, and demise will be key in understanding how phototrophic blooms impact ecosystem function and biogeochemical cycling.

## Materials & Methods

### Study site and sample collection

Trunk River Lagoon (TRL) estuary (41°32’08.4“N 70°38’26.6”W) is a brackish coastal lagoon on Cape Cod located between Oyster Pond, a kettle pond with a salinity maintained between 2-4 PSU ^82^, and Vineyard Sound. The kettle pond exchanges tidal water with Vineyard Sound through a culvert at the northern end of Trunk River Lagoon. Trunk River experiences periodic sedimentation from storms ^82^ and frequently has a sulfidic odor with gas bubbles emanating from the sediment. Salinity is dynamic within the lagoon and vertical salinity gradients often result in stratification, with the surface waters having salinity <10 and bottom waters reaching 30 ^5^. Seasonal GSB blooms occur from May to October in the southern region of the lagoon, where water depths range from approximately 0.1-1 m. GSB blooms form in the water column with an upper boundary of 10-30 cm below the water surface and extend to water depths of up to 80 cm, with exact depths varying based on lagoon’s bathymetry and hydrological conditions. Samples were collected from August-October 2021, where a single GSB bloom was sampled over a period of 7 consecutive days from August 26th-September 1st, and then again on September 10th, October 20th, and finally on October 21st when the bloom terminated.

Three types of samples were collected from the bloom: water, suspended biofilm-like matrix, and microbial mat (Fig. 1). Water samples of both the bloom and overlying water column above the bloom were collected in Ar-flushed, evacuated (BrandTech vacuum, 13-688-000) 1 L anaerobic media bottles (Chemglass, NC0364556). Open tubing (Masterflex, 06402-07) was lowered to the desired water depth and then connected to the evacuated bottles using a LuerLok connector and 21 G needle. Due to a vacuum pressure of approximately ∼900 mbar inside media bottles, approximately 900 mL of sample flowed into the bottle before the inflow subsided and the tubing was disconnected. Microbial mats, which developed at the sediment surface below the bloom, were collected using polystyrene weighing dishes (Fig. 1d). A suspended biofilm-like matrix that developed at the interface of the bloom and the overlying water column was sampled with a syringe (Fig. 1e-f, referred to as a “suspended biofilm”). All samples were stored on ice and transported back to the laboratory for processing.

Water samples were filtered onto 0.22 µm polyethersulfone Sterivex filters (SVGP01050) for 16S rRNA gene sequencing and metaproteomics. Additional metaproteomics samples of bloom supernatant were collected by centrifuging 100 mL of bloom at 20,000 x g for 15 minutes at 4°C. Samples for metagenomics were collected via sequential filtration using filters with pore sizes of 5 µm, 2 µm, 0.22 µm, 0.1 µm, and 0.025 µm (MCE Membrane Filters, Millepore Sigma) to generate samples of the following size fractions: > 5 µm, 2-5 µm, 0.22-2 µm, 0.1-0.22 µm, and 0.025-0.1 µm. Suspended biofilm samples were centrifuged for 5 minutes at 4°C at 20,000 x g and the supernatant discarded. Microbial mat samples were directly transferred into sterile 2 mL tubes. All samples were stored at -80°C until analysis.

### Water Column Chemistry

Macroscale measurements of pH, temperature, dissolved oxygen, salinity, and depth were made using a multi-parameter probe (YSI, Hydro lab DS5). Measurements were made adjacent to sample collection area and prior to sampling to avoid any disturbing the bloom or sediment.

For sulfide (H_2_S) analysis, 0.25 mL of sample was removed from sealed anaerobic media bottles using a needle and syringe and immediately added to 6 mL of 2 % zinc acetate. Samples were stored at 4°C until analysis with a spectrophotometric assay, as described by Cline ^83^. Briefly, samples were incubated in the dark for 30 to 45 min with 5 mL of diamine dye, and absorbance at 670 nm was read on a Shimadzu UV-VIS Spectrophotometer (UV-1900i).

To sample for dissolved inorganic nutrients, samples collected in media bottles were syringe-filtered directly through 0.22 µm polyethersulfone Sterivex filters (SVGP01050), and filtrate was collected in acid-washed 15 mL polypropylene tubes. Samples were immediately frozen and stored at −20°C until analysis by the Louisiana State University’s Wetland Biogeochemistry Analytical Services Laboratory using an O.I. Analytical Flow Solution IV autoanalyzer. The detection limits were 0.086 µM (NO*_x_*), 0.105 µM (NH_4+_), 0.046 µM (PO_43−_), 0.087 µM (Si(OH)_4_), and 0.177 µM (SO_42-_). Dissolved inorganic nitrogen (DIN) was determined as the sum of [NO*_x_*] and [NH_4+_]. Dissolved inorganic phosphorus (DIP) as [PO_43−_].

### Stable CNS Isotopes

Water samples were filtered onto pre-combusted 25 mm GF/F filters (Cytiva Whatman 1825025), with volume ranging from 9 mL (high cell density) to 71 mL (low cell density) per filter based on bloom density. Filters were frozen at -20°C until analysis. Prior to analysis, filters were dried overnight at 60°C. Bulk nitrogen (*δ*^15^N), carbon (*δ*^13^C), and sulfur (*δ*^34^S) isotope compositions were determined using a PDZ Europa 20-20 Continuous-Flow Isotope Ratio Mass Spectrometer system by the Stable Isotope Facility at the Marine Biological Laboratory (Woods Hole, MA). Stable isotope compositions are reported in delta notation using the following equation: *δ^a^*X (‰) = (*R_sample_*/*R_standard_* − 1) *x* 1000 where R is the ratio of ^15^N/^14^N, ^13^C/^12^C, or ^34^S/^32^S. Atmospheric nitrogen, Vienna Peedee Belemnite (VPDB) and Vienna-Canyon Diablo Troilite (VCDT) were used as R*_standard_* values for nitrogen, carbon, and sulfur, respectively.

### Nucleic acid Extraction

Nucleic acid extraction was performed using a chloroform-based method adapted from ^84^. Sterivex filters were cut with a sterile blade. Mat and suspended biofilm samples were weighed to approximately 0.2 g. Filters, mat, and suspended biofilm-like matrix samples were placed in 1 mL of DNA extraction buffer (DEB; 100 mM Tris-HCl (pH 8), 100 mM EDTA (pH 8), 100 mM sodium phosphate buffer (pH 8), 1.5 M NaCl, 1% CTAB). The extraction mix underwent three freeze-thaw cycles (-20°C to 37°C), followed by incubations with lysozyme (2 mg/mL f.c., 40 kU/mL) at 37°C for 30 min, and Proteinase K (0.2 mg/mL f.c., 6U/ml) at 37 °C for 30 min. Sodium dodecyl sulfate (SDS; 1% f.c.) was added and the extraction mix incubated at 65°C for 2 h. Extraction mixes were treated twice with chloroform-isoamyl alcohol (24:1, pH 8) and the aqueous supernatant was collected by centrifugation at 3,200 x g (10 min). Nucleic acids were precipitated for 2 h at RT with 100 % isopropanol (0.6:1 isopropanol:supernatant). Nucleic acids were collected by centrifugation at 20,000 x g for 30 min and washed twice with ice-cold 70 % ethanol. Pellets were briefly air-dried and then resuspended in 100-250 µL of nuclease-free water. A detailed protocol is available on protocols.io ^85^. DNA quality and quantity were assessed with a Qubit 2.0 fluorometer and NanoDrop 2000 spectrophotometer.

### 16rRNA V4V5 Amplicon Sequencing & Analysis

16S rRNA gene amplicon libraries targeting the hypervariable V4-V5 region were prepared by the Keck Sequencing Facility at the Marine Biological Laboratory using universal bacterial and archaeal primer sets: 518F 5’-CCAGCAGCYGCGGTAAN-3’ & 926R 5’-CCGTCWATTYNTTTRANT-3’, and 517F 5’-GYYTAAARNRYYYGTAGC-3’ & 958R ‘5-CCGGCGTTGANTCCAATT-3’. The following polymerase chain reaction (PCR) conditions were used for both primer sets: initial denaturation at 94°C for 3 min followed by 30 cycles of denaturation at 94°C for 30 s, annealing at 57°C for 45 s, and extension at 72°C for 1 min, followed by a final extension at 72°C for 2 min. PCRs were performed in triplicate using Platinum SuperFi DNA Polymerase (Invitrogen, 12351010). Extraction blanks and no-template blanks were used as negative controls. PCR products were visualized on an Agilent TapeStation 4200 using the D1000 ScreenTape assay (Agilent, 5067-5583 and 5067-5582). AMPure XP beads (Beckman Coulter, A63881) were used to purify and concentrate libraries, which were quantified and pooled in equimolar ratios. The 425-625 bp region of the bacterial and archaeal V4-V5 amplicons were size selected on a BluePippin system (Sage Science, BDF1510). Multiplexed amplicon pools were sequenced on an Illumina MiSeq PE250 platform (Illumina, MS-1023003), and data were demultiplexed by the sequencing facility.

Amplicon sequence data were processed in R 4.4.0 ^86^ using DADA2 (v1.32.0) ^87^. Reads containing ambiguous bases were removed (maxN=0). Primers were trimmed using cutadapt (V4.2) ^88^, and paired-end reads were filtered with a maximum expected error rate of 2 (maxEE=2), minimum quality scores (trunQ=2), and truncated based on quality plots (bacterial V4-V5: truncLen=(220,200); archaeal V4-V5:trunLen=(210,180)). Error rates were learned using learnErrors, the DADA2 sample inference algorithm was applied to dereplicated data and forward and reverse reads were merged. Two samples were reanalyzed in a subsequent sequencing run. Merged data from the two runs were combined to remove chimeras using removeBimeraDenovo (method=consensus). Amplicon sequence variants (ASVs) were assigned taxonomy in DADA2 using a Bayesian classifier with the SILVA reference database (v138)^89,90^. The average number of raw reads per sample was 143,786 ± 54,997 (n=44), and the average number of retained reads per sample after DADA2 processing and chimera removal was 103,659 ± 38,985 (n=44).

For bacterial 16S rRNA gene sequence analysis, blank extraction controls were used to check for contamination with the prevalence method of decontam with a threshold of 0.1 ^91^. A total of eight contaminant ASVs were identified and removed. ASVs that were identified as mitochondria (n= 54) or chloroplasts (n=144) were also removed from the bacterial dataset. A total of 3,566 bacterial ASVs were identified. A total of 2,207 ASVs occurred in bloom samples (n=10), 767 ASVs in suspended biofilm-like matrix samples (n=2), 1,365 ASVs in microbial mat samples (n=4), 1,420 ASVs in surface and water samples (n=4). Chloroplast ASVs were assigned taxonomy using the PR^2^ (V5.0) eukaryotic database ^92^ and further analyzed separately from the bacterial dataset. In the archaeal dataset, no contaminants were identified using blank extraction controls and decontam (0.1 threshold) [32]. A total of 2,632 archaeal ASVs were identified in bloom samples (n=10), 971 ASVs in suspended biofilm-like matrix samples (n=2), 1,382 ASVs in microbial mat samples (n=4), and 2,665 ASVs in surface water samples (n=4). Analyses and plots were performed in R using phyloseq ^93^.

### LoopSeq full-length 16rRNA Sequencing & Analysis

LoopSeq full-length 16S rRNA gene sequences were obtained using synthetic long reads as performed by the manufacturer (Loop Genomics, Element Biosciences). Full length 16S rRNA genes were amplified using the following primers: 27F 5’-AGAGTTTGATCMTGGCTCAG-3’ & 1492R 5’-TACCTTGTTACGACTT-3’^94^. Unique molecular barcode labels were attached to each individual 16S rRNA gene sequence, which allows gene fragments to be sequenced and reconstructed into full length 16S rRNA gene sequences. Libraries were sequenced on an Illumina platform using 2 x 150 bp paired-end chemistry. LoopSeq data were processed in in R 4.4.0 ^86^ using DADA2 (v1.32.0) ^87^. To select for complete, full-length 16S rRNA gene sequences, the function removePrimers was used with an allowed mismatch of 2 basepairs, and sequences lacking both forward and reverse primers were discarded. Remaining sequences were filtered using the following parameters: minimum length of 1000 bp (minLen = 1000), maximum length of 2000 bp (maxLen = 2000), minimum quality score (minQ = 3), maximum expected error rate of 1 (maxEE = 1), and no ambiguous bases (maxN = 0). Reads were dereplicated with derepFastq, error rates were learned using learnErrors with a bandsize of 32 (BAND_SIZE = 32) and the error estimate function was set to the PacBio Error Function (errorEstimationFunction=PacBioErrfun). The DADA2 algorithm was run with the p-value threshold set to 1^−10^ (OMEGA_A = 1e-10), a bandsize of 32 (BAND_SIZE = 32), and single detection (DETECT_SINGLETONS = TRUE). Chimeras were removed using removeBimeraDenovo (method = consensus) and taxonomy was assigned using assignTaxonomy with the SILVA reference database (v138) ^89,90^. The average number of reads was 12,494 ± 6,669 per sample.

To distinguish between short-read and LoopSeq full-length ASVs, those generated from LoopSeq are referred to as L-ASVs. L-ASVs that classified as mitochondria (n = 52) or chloroplasts (n = 317) were removed from the bacterial LoopSeq dataset. A total of 3,757 LoopSeq full-length bacterial 16S rRNA gene amplicon sequence variants (L-ASVs) occurred in bloom samples (n = 13), 747 L-ASVs in suspended biofilm-like matrix samples (n = 2), 1,177 L-ASVs in microbial mat samples (n = 3), and 2,294 L-ASVs in surface water samples (n = 5).

A phylogeny was constructed using the eight most abundant L-ASVs from green sulfur bacteria present in the bloom (genus *Prosthecochloris*), 19 *Prosthecochloris* and 2 *Chlorobium* reference sequences from NCBI GenBank. Sequences were aligned using MUSCLE^33^ V5.1 and a maximum-likelihood tree was constructed using Geneious Tree Builder (2.1.11) in GeneiousPrime (2023.0.1). The phylogeny was visualized using the Interactive Tree of Life (iTOL)^95^.

### Shotgun Metagenomic Sequencing & Analysis

Shotgun metagenomes were sequenced by Psomagen, Inc. Briefly, initial DNA quality was assessed using the Picogreen assay (Thermo Fisher) and the genomic DNA ScreenTape assay (Agilent). Libraries were prepared using the Illumina DNA Prep Kit by tagmenting DNA with bead-linked transposomes at 55°C for 15 minutes to fragment and tag DNA with adapter sequences. Tagmented samples were PCR amplified using a limited-cycle PCR program with indexes from the Illumina Nextera DNA Unique Dual Indexes kit. PCR products were purified with magnetic beads and the D5000 ScreenTape assay (Agilent) and PicoGreen assay (Thermo Fisher) were used to check quality. Highly concentrated libraries were normalized to 5 nM. Approximately 1.5-2.0 nM of libraries were loaded onto the flow cell and sequenced on an Illumina NovaSeq6000 S4 (v1.5) platform with PE150 chemistry (paired-end sequencing, 150 bp).

Illumina adapters were removed using Trimmomatic (v0.36) for paired-end reads (ILLUMINACLIP:2:30:10, LEADING:3, TRAILING:3, SLIDINGWINDOW:4:15, MINLEN:65). Removal of PhiX control reads was verified by mapping reads against phiX174 (NCBI NC_001422)^96,97^ using Bowtie2 ^97^. Low-quality reads were removed using iu-filter-quality-minoche from illumina-utils (v2.12)^98^. FastQC (v0.11.4) was used to validate adapter removal and perform quality control ^99^. Reads were assembled using MEGAHIT (v1.2.9) ^100,101^, and anvi-script-reformat-fasta was used to remove contigs shorter than 1000 bp^102^, resulting in 403,187 contigs and an N50 of 2,835 in bloom samples (n=21). The metaWRAP binning module ^103^ was used to generate metagenomic bins, also known as metagenome-assembled genomes (MAGs), from contigs using MaxBin2 (v2.2.7)^104^, MetaBAT2 (v2.12.1) ^105^, and CONCOCT (v1.0.0) ^106^. Assembly indexing and read mapping were performed within metaWRAP using BWA^107^, and the resulting sample-to-assembly mapping files were provided as input into a fourth binner, SemiBin (v1.4.0)^108^, which uses a deep siamese neural network. DASTool (v1.1.6) ^109^ was used to find the optimized, non-redundant set of bins from the combined binning output from all four binning algorithms. Bin quality and completion statistics were assessed with CheckM2 predict (v1.0.1) ^110^, taxonomy was assigned with GTDB-tk classify_wf (v 2.3.0)^111^, and coverage of both contigs and bins was determined with CoverM contig and CoverM genome (v0.6.1)^112^. Proteins were annotated with prokka (v1.13) ^113^ and eggNOG-mapper (v2.1.12) ^114^. Anvi’o tools (anvi-gen-contigsdatabase, anvi-run-hmms, anvi-run-ncbi-cogs, anvi-profile) were used to build profile databases, visualize bins, and refine select metagenomic bins (anvi-refine) following anvi’o workflows ^102,115^. A phylogeny of *Prosthecochloris* MAGs from this study and published *Prosthecochloris* genomes and metagenomes was calculated with the GTDB-Tk *de novo* workflow (de_novo_wf)^111^using the bacterial marker set with *Pelodictyon phaeoclathratiforme* (NC_011060.1) as an outgroup. Tree visualization was done using the Interactive Tree of Life (iTOL) ^95^.

The predominant *Prosthecochloris* MAG (GSB-TRL01) from the above analysis was aligned against a *Prosthecochloris* contig (contig1) generated from Oxford Nanopore Technologies (ONT) long-read sequencing of a sample from the same bloom, collected on Day 8 after the bloom onset. Details of long-read sequencing and assembly are described in Weed et al. 2026 (bioRxiv). *Prosthecochloris* contig1 has a length of 2,215,208 bp and estimated completion of 99.99 % (CheckM2). Alignment of the *Prosthecochloris* GSB-TRL01 MAG with ONT contig1 was done in Geneious Prime using Mauve ^116^ and used for visualization of the proteome of GSB-TRL01, described below. Gene cluster visualization and comparison with genomes of *Prosthecochloris* CB11 and CIB 2401 was done using clinker ^117^.

### Cell lysis, sample cleanup, and metaproteomics sample preparation

Protein was extracted from bloom samples collected on Sterivex filters (intracellular proteins), from supernatant of centrifuged bloom (extracellular proteins), microbial mats, and suspended biofilm samples. From filters, biomass was extracted and lysed using a method adapted from Solis et al. ^118^. Thawed filters were divided into ∼ 1 cm^2^ pieces with a sterile scalpel, placed in 1.7 mL of lysis buffer (4% (w/v) SDS, 100 mM Tris-HCl pH 8.0), heated at 97°C for 15 minutes, incubated for 1 hour at room temperature, and briefly vortexed for 1 minute. Samples were then centrifuged at 10,000 x g for 5 minutes and supernatant was collected. The filter lysis was repeated, and the supernatants were combined. To extract protein from supernatant of centrifuged bloom samples, ∼ 12 mL of supernatant was concentrated using Amicon Ultra-15 (10 kDa cutoff; Millipore Sigma, UFC9010) filtration units by centrifuging at 4,000 x g for 30 minutes at 4°C to a volume of 300-400 µL. Lysis buffer was added at a 1:2 ratio (sample:lysis buffer) and samples were heated at 95°C for 10 minutes to generate the final lysate. Lastly, protein was extracted from microbial mat and suspended biofilm-like matrix samples by loading the sample into tubes with Lysing Matrix E (MP Biomedicals, 2 mL tube, SKU: 116914050-CF), suspending samples in SDT-lysis buffer (4% (w/v) SDS, 100 mM Tris-HCl pH 7.6, 0.1 M dithiothreitol (DTT)), and performing bead beating (3 cycles of 6 m/s for 45 s, 30 s dwell). Biofilm and mat samples were then heated at 95°C for 10 minutes and the resulting supernatant was used as the final lysate.

An overnight trichloroacetic acid (TCA) precipitation was used to remove interfering environmental compounds (e.g., humic acids) from all samples ^119^. Samples were suspended in 10% TCA and kept at -80°C overnight, then thawed and centrifuged at 21,000 x g for 15 minutes at 4°C to collect the precipitated protein. The resulting pellet was washed in 1 mL of ice-cold 100 % acetone, vortexed briefly and centrifuged at 21,000 x g for 15 minutes at 4°C. Pellets were air dried and resuspended in 60-120 µL of SDT-lysis buffer.

Samples were prepared for LC-MS/MS analysis using a modified filter-aided sample preparation method ^120^. Samples in SDT-lysis buffer were heated at 95°C for 10 minutes before being loaded onto 10 kDa PES membrane centrifugal filters (VWR, cat: 82031-348) with UA buffer (8 M urea, 0.1 M Tris-HCl pH 8.5) at a ratio of 60 µL of sample lysate to 400 µL of UA buffer. Filter units were centrifuged at 14,000 x g for 20 minutes and then washed by adding 200 μL UA buffer and centrifuging as before. 100 μL of IAA solution (0.05 M iodoacetamide in UA solution) was added to filters and mixed at 600 rpm for 1 minute (Benchmark Scientific Inc., model: H5000-HC) which were then incubated at 22°C for 20 minutes followed by washing three times with 100 μL UA before performing a buffer exchange to 50 mM ammonium bicarbonate (ABC buffer) with three washes of 100 μL ABC buffer. Proteins were digested by adding 0.8 to 1 μg of MS grade trypsin (Thermo Scientific Pierce) in 40 μL of ABC buffer to the filter units, mixing at 600 rpm for 1 minute, and incubated overnight at 37°C in a wet chamber. After, peptides were eluted by centrifuging filter units at 14,000 x g for 20 minutes, followed by an additional elution with 50 μL of 0.5 M NaCl, mixing at 600 rpm for 1 minute, and centrifugation as before. Peptide concentrations of final eluates were determined using the Pierce Micro BCA assay (Thermo Scientific Pierce) following the manufacturer’s instructions.

### 1D nanoflow liquid chromatography and tandem mass spectrometry (LC-MS/MS)

For each sample 2000 ng of peptide were separated using an UltiMate 3000 RSLCnano UHPLC system (Thermo Fisher Scientific). Samples were first loaded onto a 5 mm, 500 μm C18 Acclaim PepMap100 pre-column (Thermo Fisher Scientific) for desalting, prior to separation on a 75 μm x 75 cm EASY-spray column packed with PepMap RSLC C18, 2 μm material (Thermo Fisher Scientific) at 60°C. Peptides were separated using a 140-minute reverse-phase gradient ^121^. The eluting peptides were ionized via electrospray ionization (ESI) with an Easy-Spray source and measured using an Orbitrap Exploris 480 Mass Spectrometer (Thermo Fisher Scientific) by a data dependent acquisition ^121^. In brief, the precursor scans were acquired for a 380 to 1,600 m/z window at a resolution of 60,000 with a maximum injection time of 200 ms and a normalized AGC target of 3e6. The 15 most abundant ions were isolated for MS2 analysis (Top15), using a dynamic exclusion of 25 s. MS2 spectra were acquired at a maximum injection time of 50 ms and an AGC target of 1e5. Isolated ions were subjected to a normalized collision energy of 27 % in the HCD cell prior to measurement in the Orbitrap mass analyzer at a resolution of 15,000. An average of 81,000 spectra collected per sample.

### Metaproteomic database construction and protein identification

A protein sequence database was created using predicted protein sequences from the MAGs associated with bloom, mat, and biofilm-like web samples, as well as protein sequences predicted from unbinned contigs. We constructed this database following previous recommendations^122^, reducing redundancy using CD-HIT for sequence clustering, and prioritizing taxonomically resolved sequences from MAGs over sequences from unbinned contigs with CD-HIT-2D ^123,124^. The cRAP database (https://www.thegpm.org/crap/) was used to account for the presence of common laboratory contaminants.

We identified peptides and inferred proteins from this data by searching the MS2 spectra against the custom database using Proteome Discoverer version 2.3 (Thermo Fisher Scientific) using a previously described method ^125^. Proteins were quantified using peptide spectral counts (PSMs) and only proteins that had a false discovery rate (FDR) of less than 5 % that were designated as master proteins were retained. Spectral count data were used to calculate normalized spectral abundance factor percentages (NSAF), which is the spectral abundance factor—the spectral counts for a protein divided by amino acid length of the protein—normalized against the sum of all proteins to allow for comparison across samples and accounts for the fact that longer proteins yield more spectra. Organism-normalized NSAF were also obtained by normalizing values to the primary organism in the bloom (GSB-TRL01-NSAF) ^37,126,127^. MAG biomass contributions were determined following Kleiner et al. ^37^.

### Metaproteomic Analysis

Three different metaproteomics subsets of the data were used in analysis: (1) MAG biomass contributions, (2) normalized spectral abundance factor (NSAF) protein abundance of the entire community, and (3) GSB-TRL01 organism normalized NSAF-based protein abundance (GSB-TRL01-NSAF). Biomass data was median normalized and plotted using phyloseq and microViz packages in R ^93,128^. Multiple taxonomic rankings are provided in instances where the taxonomic resolution of the protein could not be resolved due to shared proteins between MAGs.

From the community NSAF dataset, proteins involved in oxidative and reductive sulfur pathways were selected based on their annotations from Prokka ^113^ including: sulfide quinone reductase (sqr), flavocytochrome sulfide dehydrogenase (fcc), sulfur oxidizing-sox complex (sox), dissimilatory sulfate reduction (dsr), adenylylsuflate reductase (apr), sulfur dioxygenase (sdo), sulfate adenylyltranferase (sat), and polysulfide reductase/thiosulfate reductase (psr/phs), as well as the nitrogenase protein subunits nifHDK. Phylogenetic analyses were done with published reference sequences for respective functional proteins in Geneious Prime using MAFFT Alignment and FastTree. Phylogenies were then used to determine if dsr, apr, and sat proteins were in the oxidative or reductive forms, and to determine cluster information for all sqr and nifH proteins found within the bloom ^52,129^. NSAF-abundances of sulfur cycling proteins and their associated MAG were plotted in R using phyloseq and microViz packages.

Using the GSB-TRL01-normalized NSAF dataset and the aligned GSB-TRL01 MAG, protein expression was visualized across the timeseries sampling period using anvi’o (anvi-interactive) in manual mode ^115^. The top 250 most abundant proteins from GSB-TRL01 were annotated using additional protein databases: Mantis, GhostKoala, eggNOG ^114,130,131^. Broad protein functional categories groups were determined manually and visualized with microViz.

### Total Cell Counts and CARD-FISH

Bloom samples collected in anaerobic media bottles were fixed in paraformaldehyde (2 % f.c. in 1 x PBS) for 1 h at RT, filtered onto 0.2 µm polycarbonate filters (MilliporeSigma, GTTP04700) with a 0.45 µm support filter (Sartorius, 14555400), and rinsed with excess 1 x PBS. For cell counts, filter segments were incubated with DAPI (4’,6-diamidino-2-phenylindole, 1 µg mL^−1^) for 10 minutes at RT, rinsed with ultrapure water and with 80 % ethanol, and air dried.

Catalyzed reporter deposition fluorescence in situ hybridization (CARD-FISH) was performed following Pernthaler et al.^132^ with modifications using HRP-probes (biomers.net) targeting green sulfur bacteria (GSB-532)^133,134^ and a nonsense, negative control probe (NON338)^135^. Cells were permeabilized using a lysozyme solution of 1000 kU mL^−1^ in 0.05 M EDTA [pH 8.0], 0.1 M Tris-HCl [pH 7.5], incubated for 1 h at 37°C, and washed in ultrapure water. Endogenous peroxidases were inactivated by incubating filters with 0.01 M HCl for 15 minutes at RT, followed by washing with ultrapure water and 96 % ethanol (EtOH). Filters were then air dried prior to hybridization with HRP-labeled probes. A 10 % w/v sterile blocking solution was prepared with blocking reagent (Roche, 11096176001) in maleic acid buffer (100 mM maleic acid and 150 mM NaCl [pH=7.5]). 300 µL of hybridization buffer (0.9 M NaCl, 20 mM Tris-HCl [pH 7.5], 10% w/v dextran sulfate, 0.02 % w/v sodium dodecyl sulfate (SDS), 1 % w/v blocking reagent, and 10 % (GSB-532) or 35 % formamide (NON-388) were combined with 1 µL of HRP-probe working solution (50 ng µL^−1^; 0.16 ng µL^−1^ final), and filters were incubated at 46°C for 2-3 h. Filters were washed with buffer (0.45 M (GSB-532) or 0.08 M NaCl (NON-388), 5 mM EDTA [pH 8.0], 20 mM Tris-HCl [pH 7.5], 0.01 % w/v SDS) for 10 minutes at 48°C and in 1 x PBS for 15 minutes at RT. Filters were then transferred to amplification buffer (2 M NaCl, 0.1 % w/v blocking reagent, 0.0015 % H_2_O_2_, 10 % w/v dextran sulfate; in 1 x PBS) containing Alexa Flour 488 (Invitrogen, B40957) tyramide (1 mg mL^−1^) in a ratio of 1:500 and incubated in the dark for 30 minutes at 46°C. Filters were incubated in 1 x PBS for 10 minutes at room temperature in the dark, followed by washing in ultrapure water and 96% EtOH. A detailed protocol is available on protocols.io^136^.

Filters for total cell counts (DAPI-stained) and GSB-specific cell counts were then embedded in an antifade mounting medium (Citifluor:Vectashield, 4:1) on a microscopy slide, and cells were enumerated using a counting grid (100 x 100 µm) and 20 independent fields of view per sample on a Zeiss Axiolab epifluorescence microscope.

## Data Availability

The following can be found in an Open Science Framework supplemental data repository (https://osf.io/skuq7/overview?view_only=4ecb92165508457e8fdad89780139af3): a list of samples and corresponding analysis performed on each sample; dissolved inorganic nutrient data; stable isotope data; YSI probe data; metagenomic bins with associated taxonomy and completeness information; trees, alignments, and fasta files used for all phylogenies; metaproteomic data tables and .faa protein files for each bin. Raw protein spectral data and a sequence database of all 1,253,648 protein sequences is available in the PRIDE repository (Accession: PXD064785 [REVIEWER TOKEN: ylrzohYwCArL (https://www.ebi.ac.uk/pride/login)]). All raw sequencing data can be found on the NCBI Sequence Read Archive (BioProject: PRJNA1417754). Files and scripts used in amplicon, LoopSeq, metagenomic and metaproteomic analysis and to generate figures are available at https://github.com/moyn413/TrunkRiverChlorobi.

## Acknowledgements

This work was funded by The Simons Foundation (Award: 824763, S.E.R), The US National Science Foundation (PRFP #2205993 to M.A.M and IOS #2426305 to M.K.), and an NSF-ACCESS computing award (BIO220128, M.A.M). LC-MS/MS measurements were made in the Molecular Education, Technology, and Research Innovation Center (METRIC) at North Carolina State University, which is supported by the State of North Carolina, USA. The authors thank Marshall Otter for use of his canoe and assistance with isotope analysis, Katie Haviland and Samuel Kelsey for Cline assay training, Bailey Fallon and the MBL Keck Facility for amplicon sequencing, Joe Vineis and the EBAME course for help and inspiration using Anvi’o, and Rich Fox for support using the JBPC server at MBL.

