## Supplemental Figures for "Anoxygenic phototrophic Chlorobi use broad metabolic and resource acquisition strategies to support stable near-clonal blooms"

#### Contents

**1 Sulfate and Sulfide Concentrations 2**

**2 Bloom vs Surface Water Silica, Nitrate, and Nitrite Concentrations 3**

**3 Timeseries of Dissolved Inorganic Nutrients in Bloom 4**

**4 YSI Probe Data 5**

**5 Microbial Mat and Suspended Biofilm-like Matrix Microbial Community Composition 6**

**6 Metagenome-assembled Genome (MAG) abundances 7**

**7 Archaeal Community Composition of Bloom 8**

**8 Sulfide quinone reductase (Sqr) Phylogeny 9**

**9 Polysulfide/thiosulfate reductase (psrA/phsA) Phylogeny 10**

**10 Microbial Mat and Suspended Biofilm-Like Matrix Sulfur Cycling Protein Abundances 11**

**11 Nitrogenase (nifH) Phylogeny 12**

**12 Nitrogenase (nifDHK) Protein Abundances 13**

### 1 Sulfate and Sulfide Concentrations

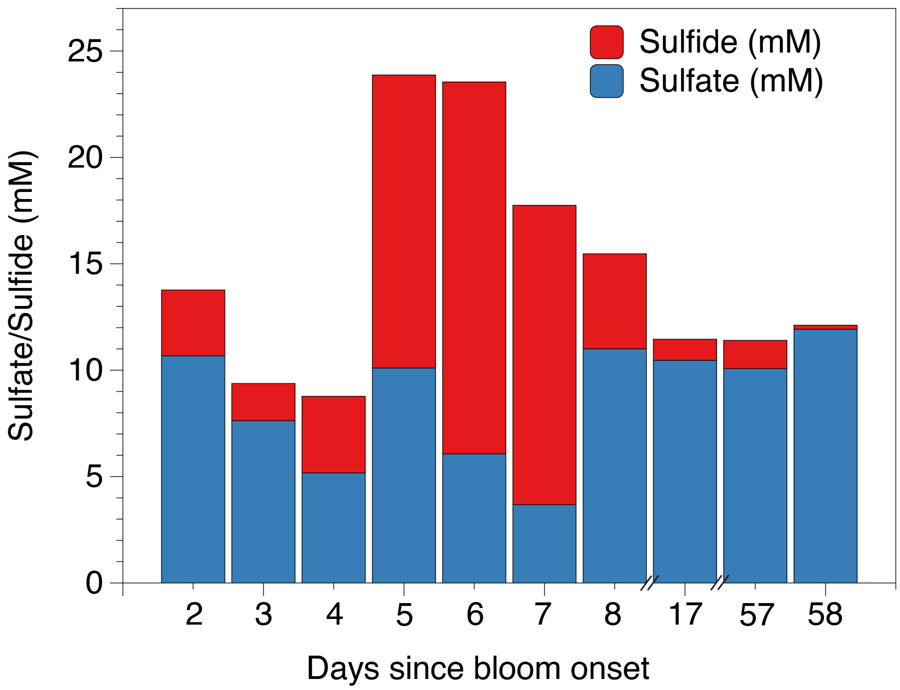


Figure S1: Stacked bar plots of the sulfate (mM) and sulfide (mM) concentrations found within GSB bloom samples from 26-Aug-2021 (Day 2) to 21-Oct-2021 (Day 58). Samples were collected at approximately 0.4m depth within Trunk River Lagoon estuary.

#### 2 Bloom vs Surface Water Silica, Nitrate, and Nitrite Concentrations

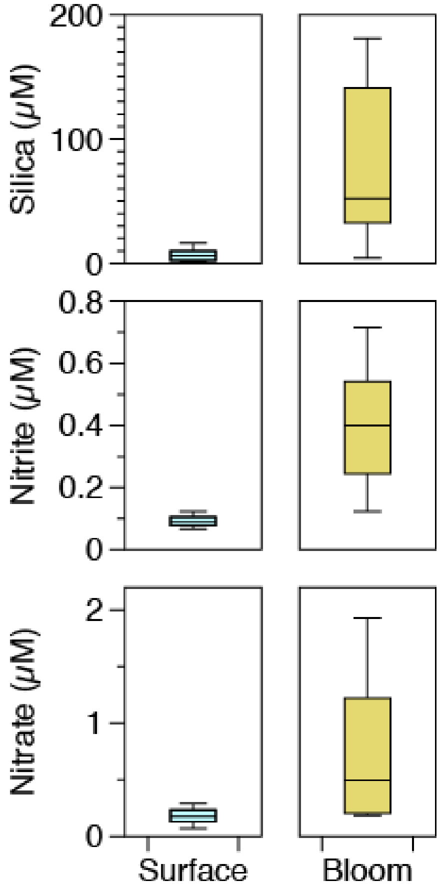


Figure S2: Silica (μM), nitrate (μM), and nitrite (μM) concentrations between bloom and overlying surface water averaged over the bloom period (Day 2 to Day 58) sampled at approximately 0.4 m depth in Trunk River Lagoon estuary.

#### 3 Timeseries of Dissolved Inorganic Nutrients in Bloom

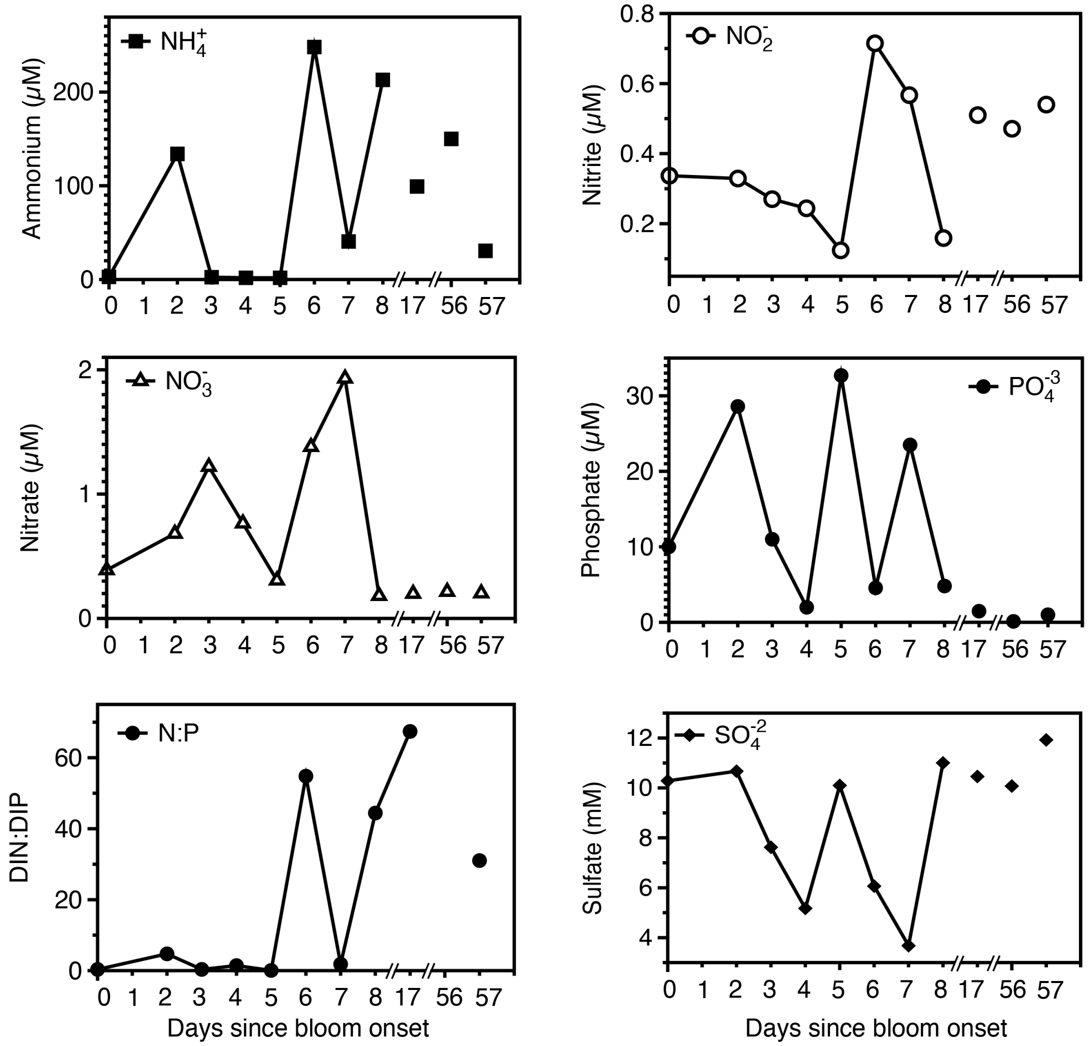


Figure S3: Dissolved inorganic nutrient data from within the GSB bloom over the period of 24-Aug-2021 (Day 0) to 21-Oct-2021 (Day 58). Samples were collected at approximately 0.4 m depth within Trunk River Lagoon estuary. Dissolved inorganic nitrogen (DIN) was determined as the sum of ammonium, nitrate, and nitrite, and dissolved inorganic phosphorus (DIP) is the concentration of phosphate. Note that the DIN:DIP ratio of Day 55 was elevated (1165.8) and excluded from the plot. Raw data is available in the supplementary data repository.

### 4 YSI Probe Data

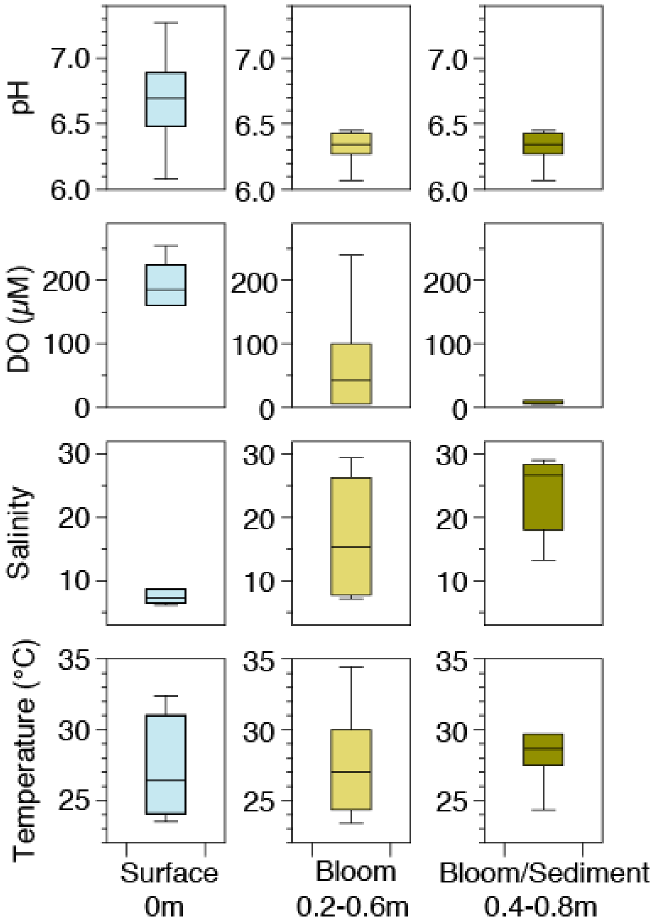


Figure S4: pH, dissolved oxygen (µM), salinity, and temperature (°C) measurements made from Day 2 to Day 7 (n=6) during the bloom period using a YSI multiparameter sonde in Trunk River Lagoon estuary.

### 5 Microbial Mat and Suspended Biofilm-like Matrix Microbial Community Composition

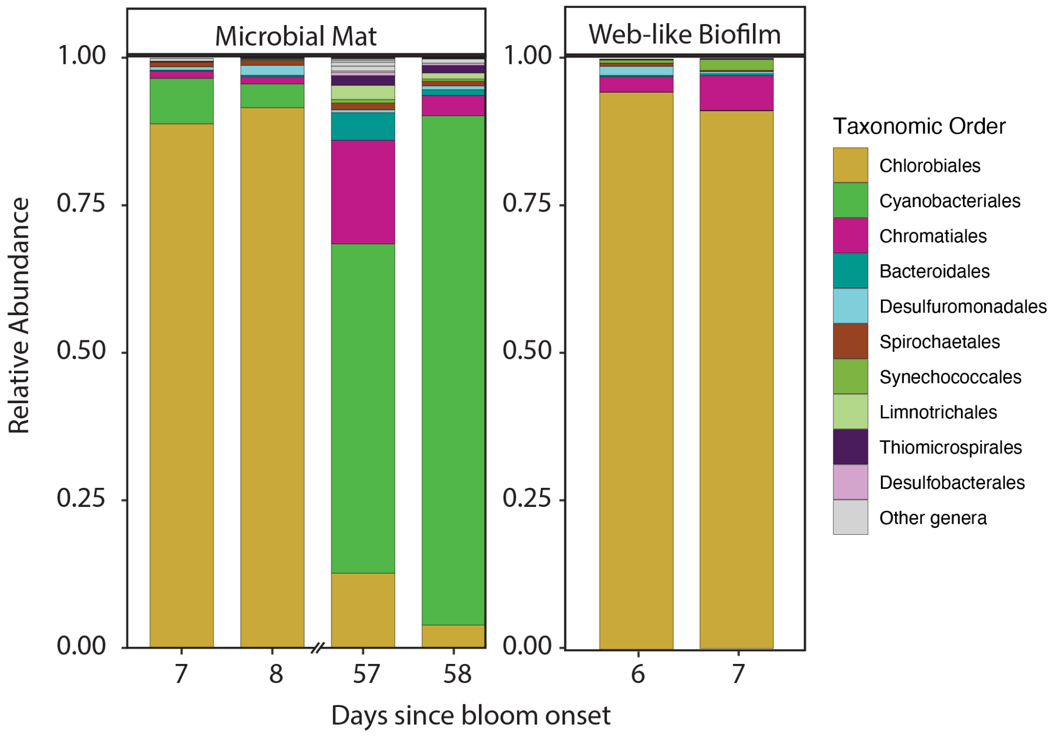


Figure S5: 16S rRNA gene-based relative abundance of the V4V5 region of microbial mat sampled below bloom and suspended biofilm-like matrix sampled at the bloom interface with the overlying water column. Sample day 6 corresponds with 31-Aug-2021 and sample day 58 with 20-Oct-2021. The legend shows the 10 most abundant taxonomic orders.

### 6 Metagenome-assembled Genome (MAG) abundances

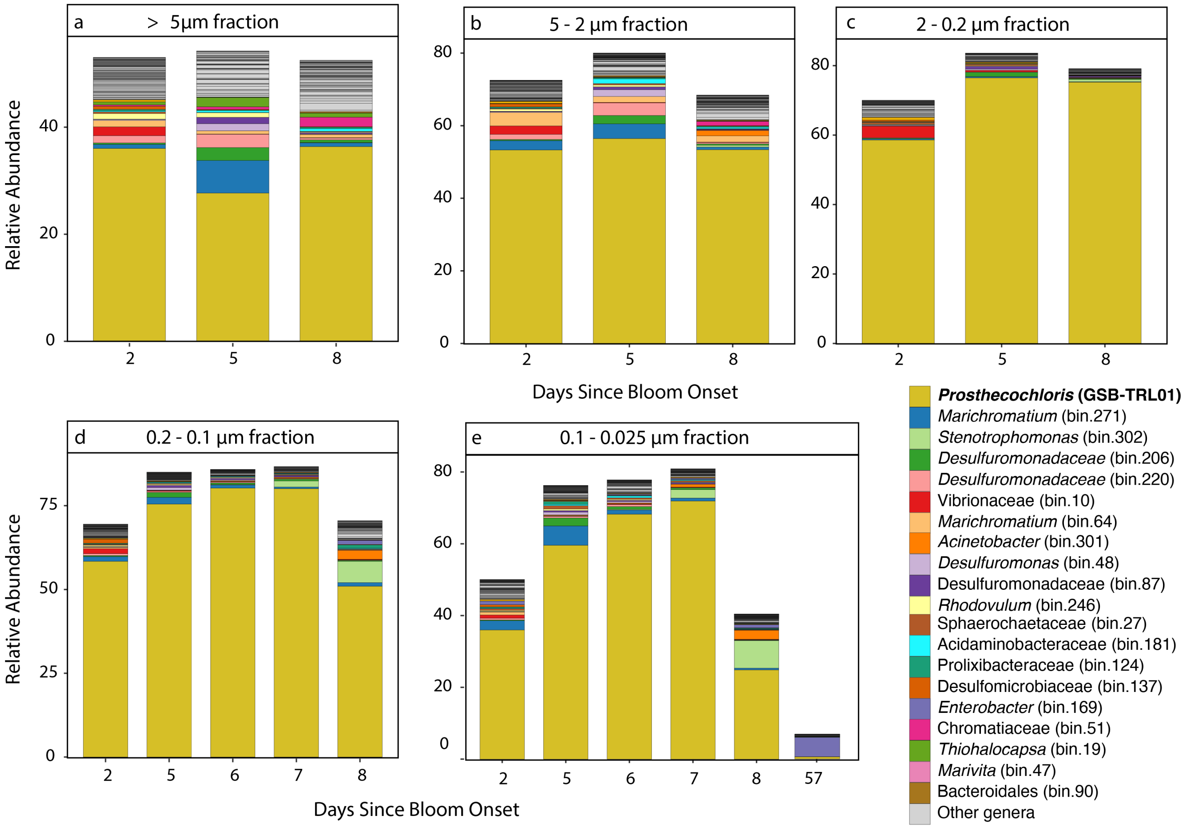


Figure S6: Relative abundances of metagenome assembled genomes (MAGs) from co-assembly of bloom samples according to the filter size fraction sequenced, (a) >5 µm, (b) 5 - 2 µm, (c) 2 – 0.2 µm, (d) 0.2 – 0.1 µm, and (e) 0.1– 0.025 µm. The top 20 most abundant MAGs are displayed. Relative abundance was determined using CoverM. Note that y-axis varies between plots. Samples were from within the GSB bloom over the period of 26-Aug-2021 (Day 2) to 1-Sept-2021 (Day 8), with Day 57 in the smallest size fraction. Samples were collected at at approximately 0.4 m depth in Trunk River Lagoon.

### 7 Archaeal Community Composition of Bloom

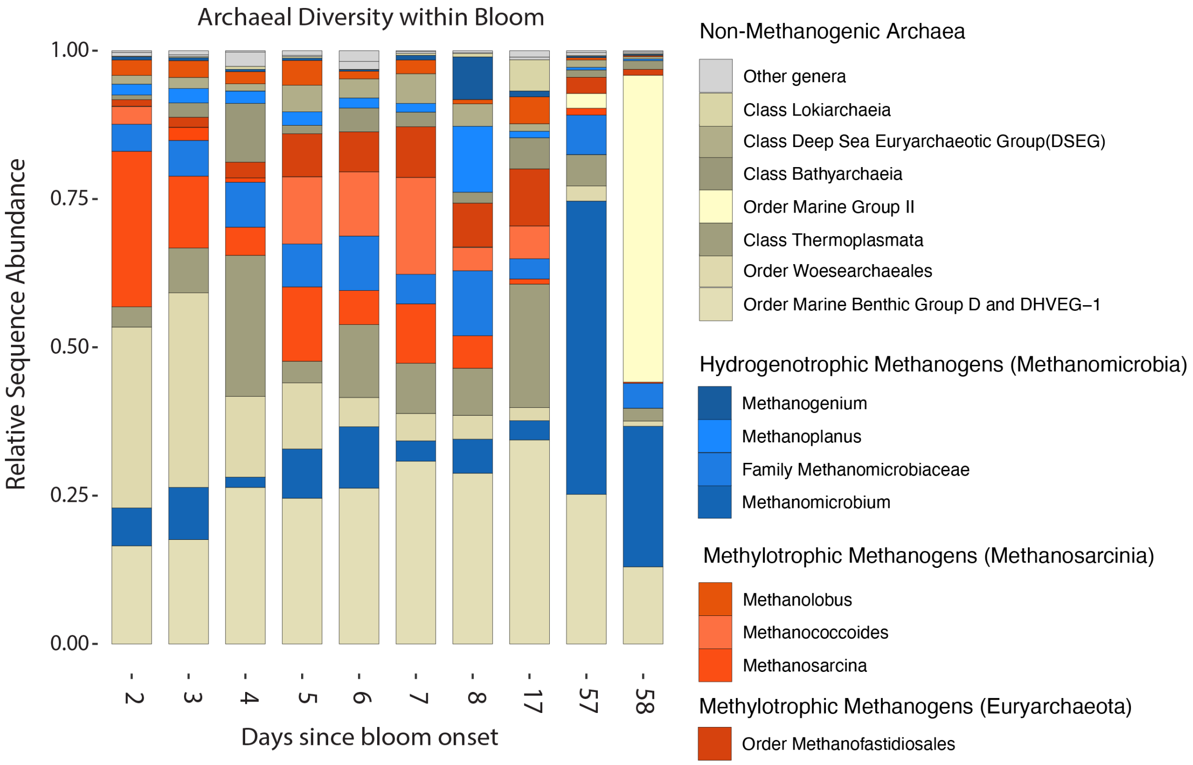


Figure S7: Relative abundance of archaeal amplicon sequence variants from short-read Illumina sequencing of the 16S rRNA gene using archaeal-specific primers (AV4V5). Samples were from within the GSB bloom over the period of 24-Aug-2021 (Day 0) to 21-Oct-2021 (Day 56) and sampled at approximately 0.4 m depth in Trunk River Lagoon. Taxonomic groups were organized based on their putative status as methanogenic vs. non-methanogenic.

### 8 Sulfide quinone reductase (Sqr) Phylogeny

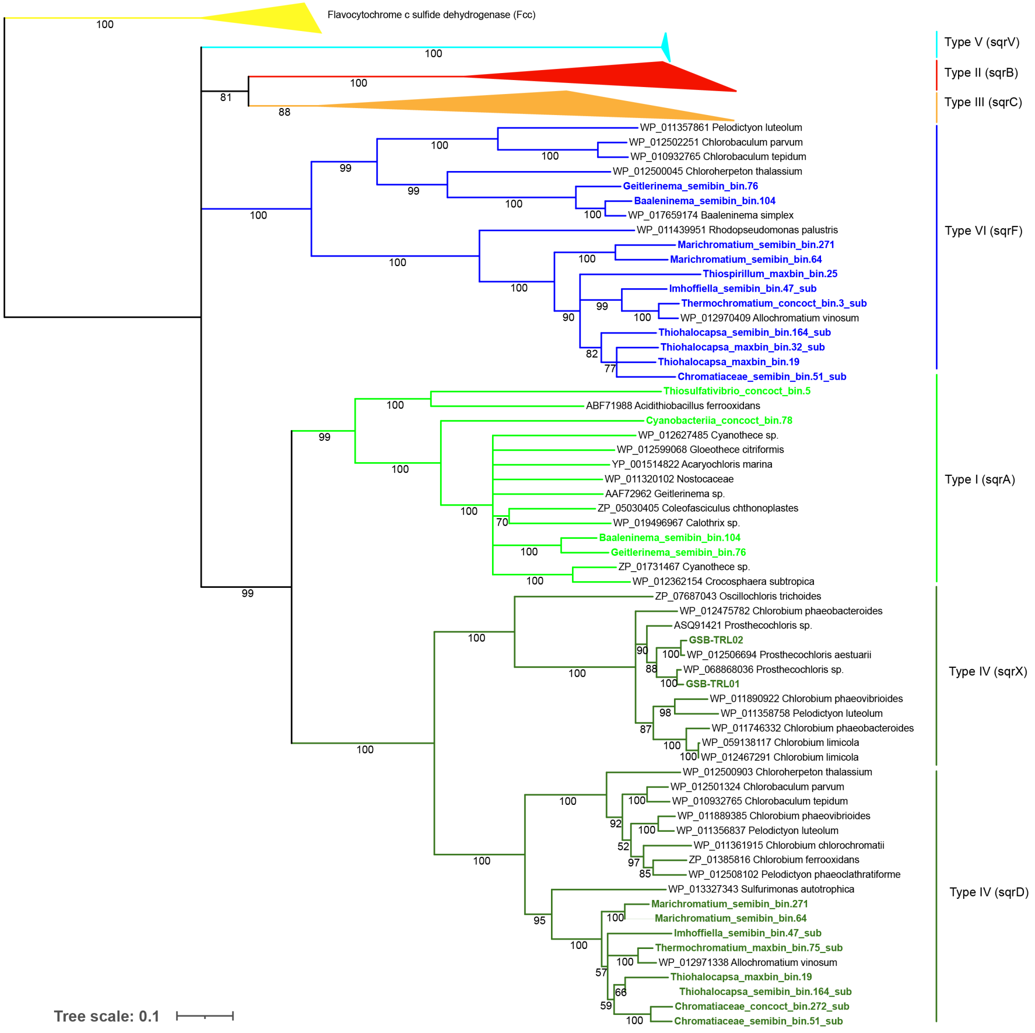


Figure S8: Sulfide quinone reductase (Sqr) phylogeny including sequences from MAGs present in Trunk River Lagoon bloom samples. Trunk River sequences and corresponding bin names are colored in blue (Type VI, sqrF) and neon green (Type I, sqrA), and dark green (Type IV, sqrX & sqrD). Type classification was modeled after Marcia et al. [1] and Grim et al. [2]. The tree file and sequences used to make the tree are available in the supplementary data repository.

#### 9 Polysulfide/thiosulfate reductase (psrA/phsA) Phylogeny

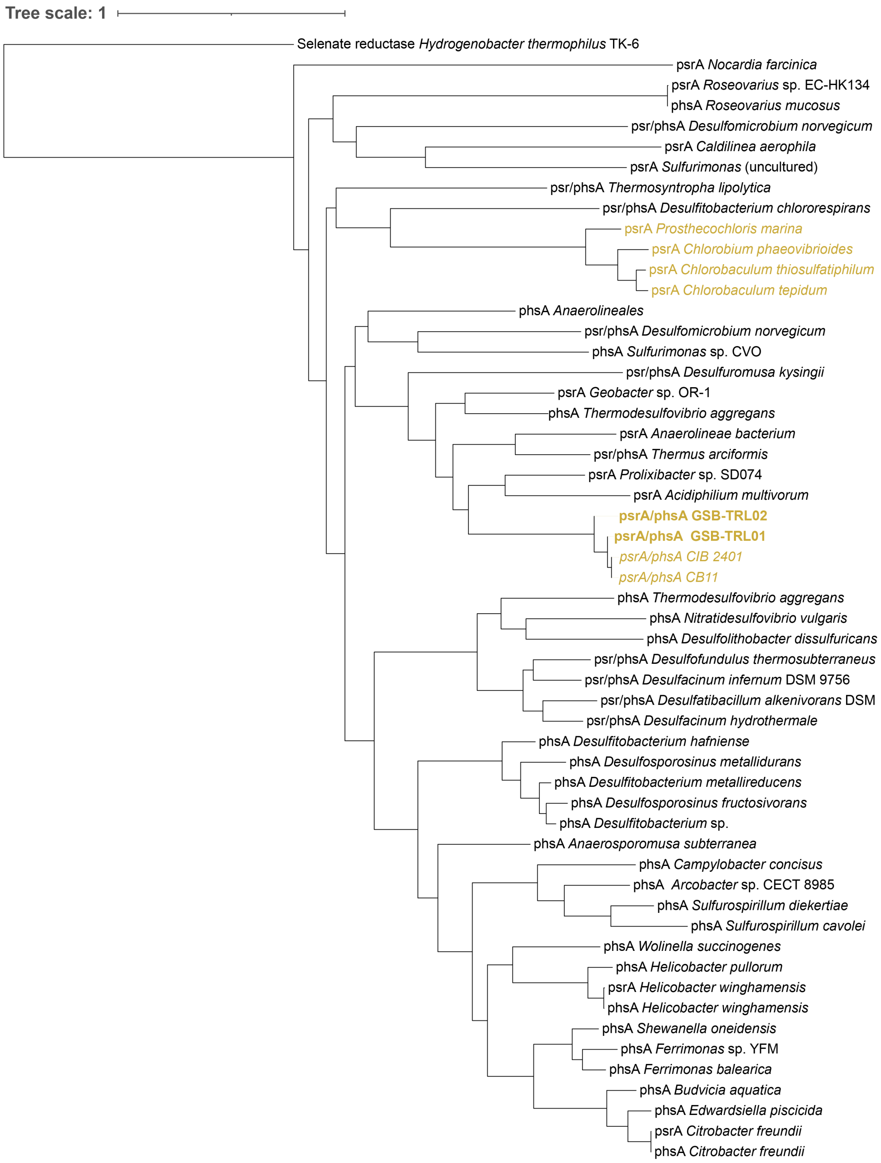


Figure S9: Phylogeny of putative polysulfide reductase/thiosulfate reductase (psrA/phsA) genes, including MAGs from this study (GSB-TRL01 and GSB-TRL02) from Trunk River Lagoon.

### 10 Microbial Mat and Suspended Biofilm-Like Matrix Sulfur Cycling Protein Abundances

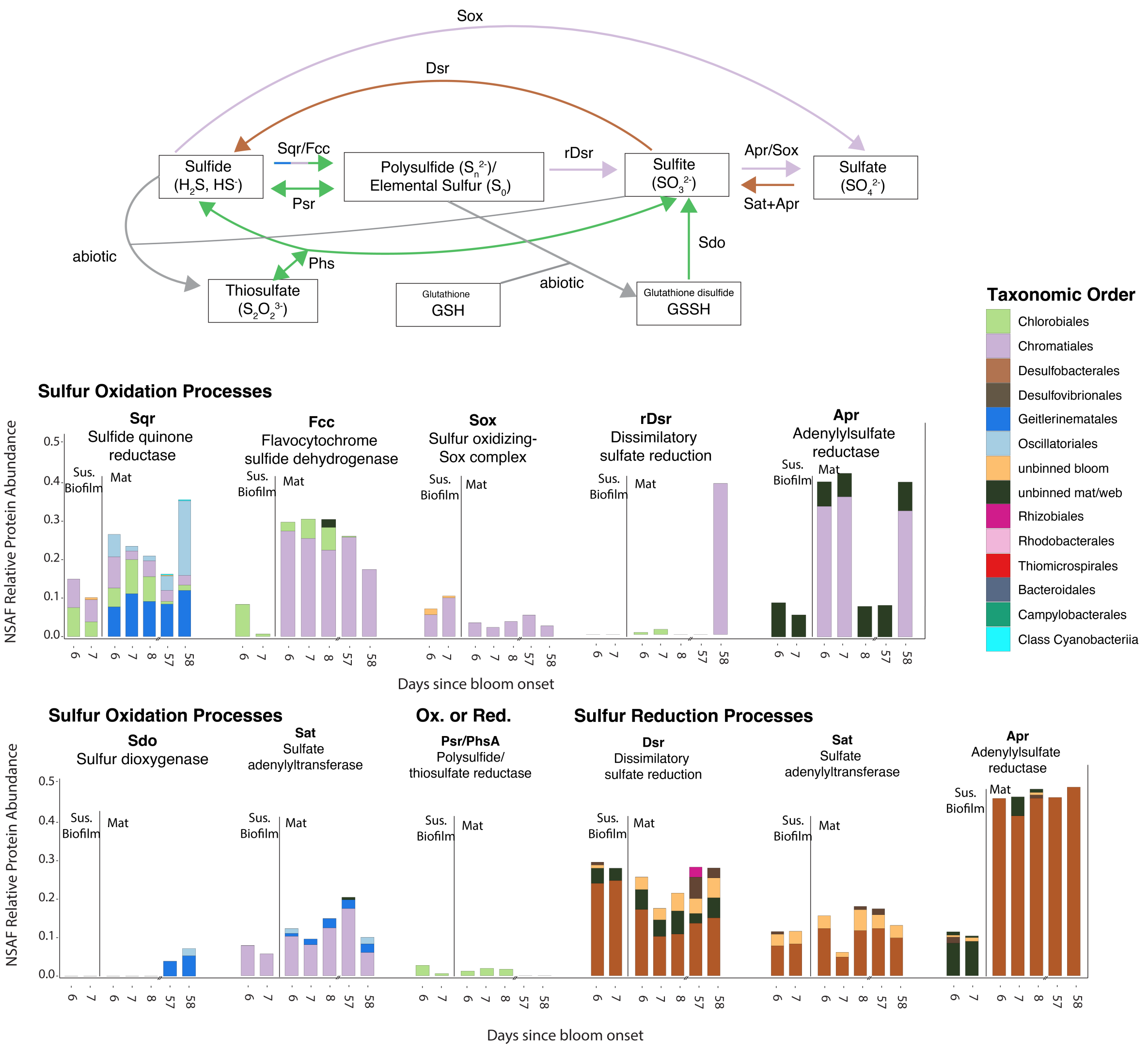


Figure S10: Normalized spectral abundance factor (NSAF) protein abundance of sulfur cycling proteins found in suspended biofilm-like matrix (Sus. Bioflm) and Mat communities the duration timeseries sampling. Sample day 6 corresponds with 31-Aug-2021 and sample day 58 with 21-Oct-2021. The legend shows the 10 most abundant taxonomic orders. Proteins are organized by oxidative vs reductive pathways and by taxonomic order based on the metagenome assembled genomes (MAGs) corresponding with each protein.

### 11 Nitrogenase (nifH) Phylogeny

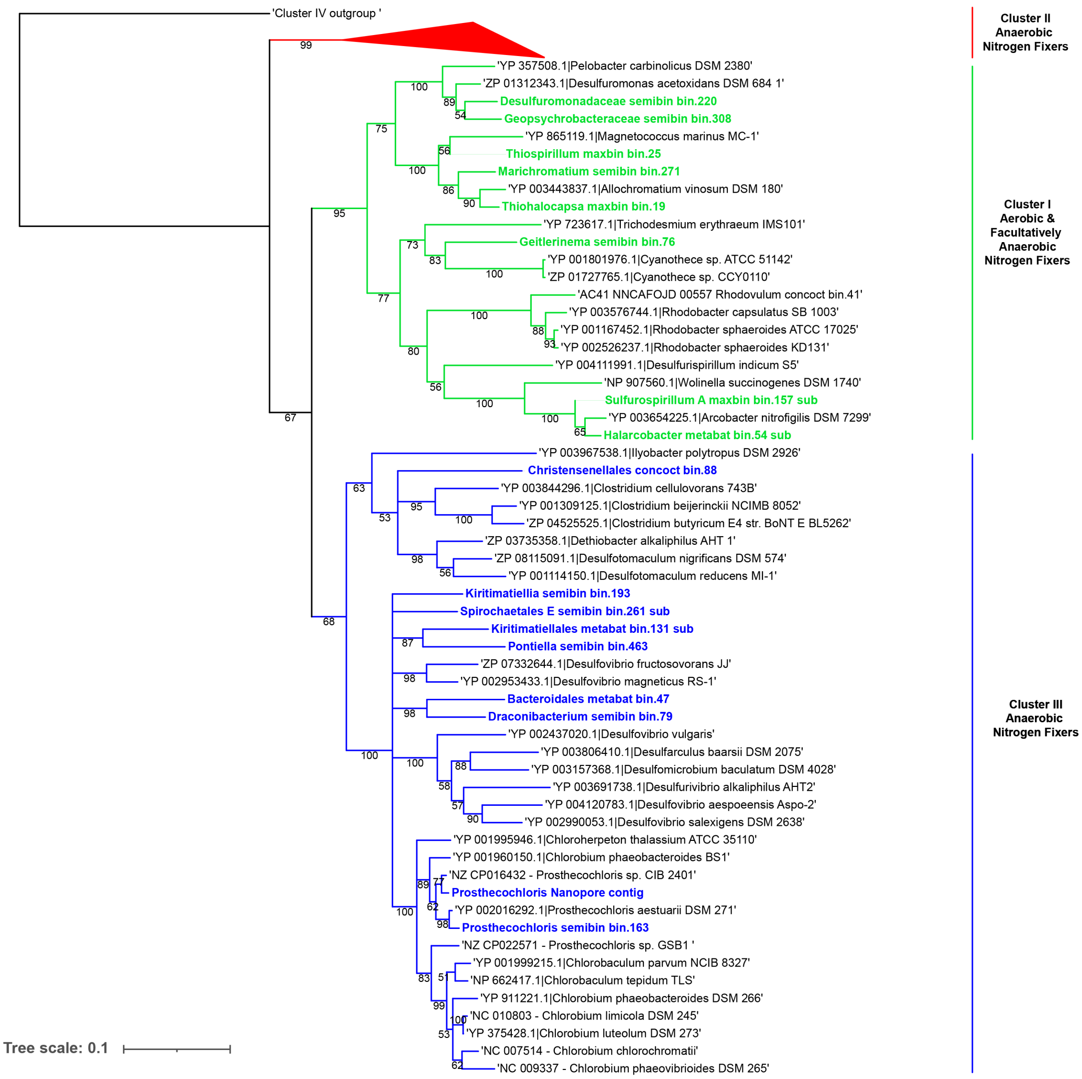


Figure S11: Nitrogenase subunit H (nifH) phylogeny including sequences from MAGs present in Trunk River Lagoon bloom samples. Trunk River sequences and corresponding bin names are colored in green (Cluster I nifH), and blue (Cluster III nifH). The phylogeny is based on selected references sequences from Moynihan et al.[3]. The tree file and sequences used to make the tree are available in the supplementary data repository.

### 12 Nitrogenase (nifDHK) Protein Abundances

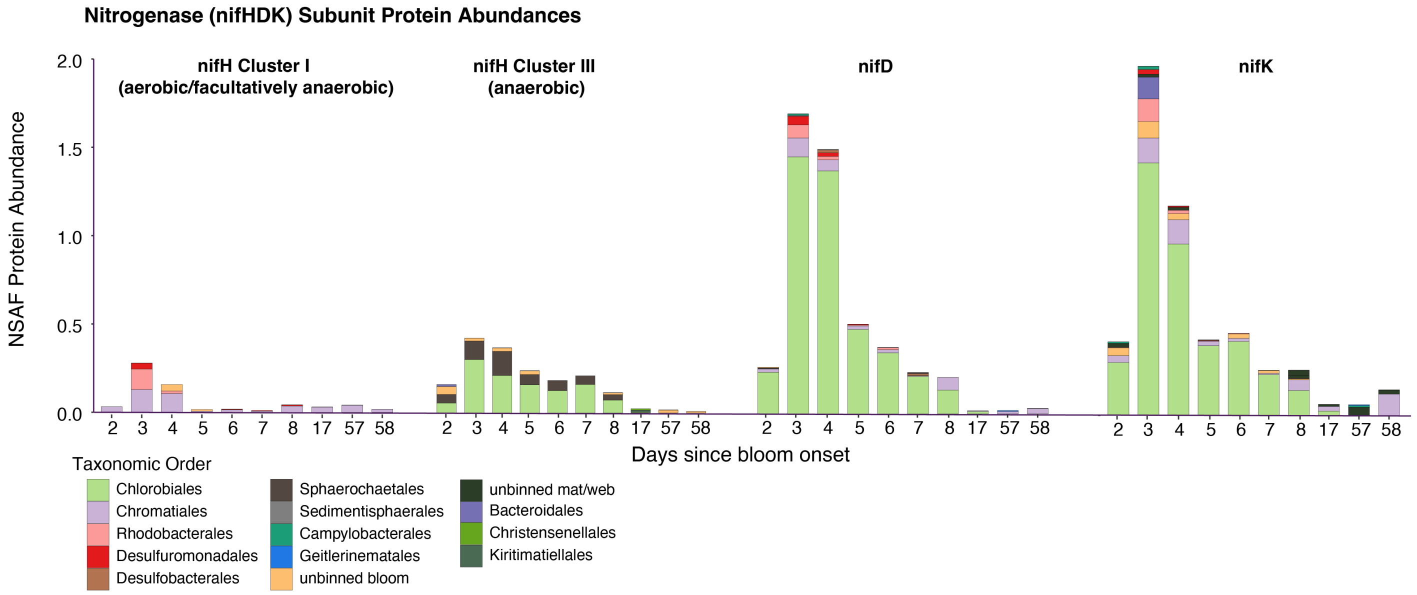


Figure S12: Normalized spectral abundance factor (NSAF) protein abundance of nitrogenase subunits nifH, nifD, and nifK found in the entire Trunk River GSB bloom community over the duration timeseries sampling (26-Aug-2021 to 21-Oct-2021). Proteins are colored based on the taxonomic order based on the metagenome assembled genomes (MAGs) corresponding with each protein. nifH proteins were separated by cluster (Clusters I & III) based on the phylogeny (Fig. S11).
